# CircEPSTI1 regulates miR-942-5p-SERPINE1-AKT1 signaling axis to enhance dengue infection and is suppressed by Tiplaxtinin

**DOI:** 10.1101/2025.05.05.652247

**Authors:** Nilanjana Das, Pandikannan Krishnamoorthy, Kushal Ramrao Junghare, Athira S Raj, Priyanka Saha, Kuhu Pal, Shouvanik Adhya, Atul Garg, Dheeraj Khetan, Ashok Kumar, Himanshu Kumar

## Abstract

Circular RNAs (circRNAs) have recently been identified as crucial regulators of gene expression in various diseases. Their role in dengue remains unexplored. In this study, circEPSTI1 induction was observed in peripheral blood mononuclear cells (PBMCs) and plasma of dengue patients. The induction of circEPSTI1 is interferon-dependent and enhances DENV infection by sponging the expression of miR-942-5p. The antiviral role of miR-942-5p is mediated by bidirectional inhibition through targeting conserved genomic sequences of the DENV genome of different subtypes across NS1, NS3, and NS5 and the host AKT1 signaling pathway. RNA-seq analysis of DENV-infected circEPSTI1 knockdown A549 cells identified SERPINE1 and AKT1 signaling pathways to be dysregulated significantly. circEPSTI1 relieves the inhibition of miR-942-5p over DENV genomic RNA and the host SERPINE1 to activate AKT1 signaling. The activation of AKT1 signaling facilitates cell survival and enhances DENV replication. The expression of SERPINE1 and circEPSTI1 was upregulated in Dengue patients, and pharmaceutical inhibition of SERPINE1 through Tiplaxtinin inhibits the DENV replication by reducing the expression of circEPSTI1. Overall, our result demonstrates the therapeutic potential of Tiplaxtinin by targeting the circ-EPSTI1-miR-942-5p-SERPINE1-AKT axis in dengue treatment.

## Introduction

Dengue is an emerging global health threat due to the increased outbreaks and its distribution to over half of the global population [1]. Dengue is linked to symptoms ranging from moderate fever in the case of Dengue fever (DF) to thrombocytopenia, vascular leakage in the case of Dengue with Warning signs (DWS), and excessive production of cytokines and organ dysfunction in the case of Severe Dengue (SD) [2]. Different serotypes of the Dengue virus (DENV1, DENV2, DENV3, and DENV4) are transmitted through *Aedes* aegypti. The DENV genome is an 11kb positive-sense RNA genome consisting of a highly conserved 5’-UTR, a lengthy ORF coding polyprotein, and a very organized, unstable 3’-UTR. Secondary infection with a different serotype can lead to severe clinical outcomes through the process known as antibody-dependent enhancement, which limits the development of DENV vaccines [3].

Host innate immunity acts as the first layer of defense against DENV infection [4]. DENV is sensed by Pathogen recognition receptors (PRRs) such as Toll-like receptors (TLR-3/7) and Retinoic-acid-like receptors (RLRs), which initiate type-I Interferons (IFN-α, IFN-β) production to limit DENV replication. RIG-I senses the DENV RNA and recruits the adaptor protein, IPS-1 (also known as MAVS, VISA, and CARDIF), subsequently activating transcription factor IFN regulatory factor 3 (IRF3) and NFκB to produce Type-I, Type-III IFN and inflammatory cytokines, respectively [5]. However, DENV utilizes multiple strategies to evade the antiviral innate immunity and establish the infection, skew physiology, and develop a range of clinical symptoms [6].

Non-coding RNAs play an important role in regulating genes and various signaling pathways that are part of the antiviral immune responses [3]. Non-coding RNAs can be categorized into small non-coding RNAs, such as microRNAs (length less than 200 nucleotides), and long non-coding RNAs, such as lncRNAs (length more than 200 nucleotides). An emerging class of transcripts, circular RNAs (circRNAs), single-stranded, covalently closed RNAs without free ends, formed through back-splicing, are shown to have higher diagnostic and therapeutic potential due to their excellent stability and presence in diverse cell types and extracellular fluids. CircRNAs were reported to regulate various biological processes in diseases like cancer, neurological disorders, and autoimmune diseases [7].

CircRNAs exert their mechanistic role by binding to RNA-binding proteins (RBPs) and are also well known to sponge microRNAs (miRNAs) [8]. miRNAs are small non-coding RNAs that post-transcriptionally regulate the expression of genes by binding to the 3’-UTR of mRNAs. miRNAs can either bind to the mRNAs of the innate immune genes or bind to the viral genomes of RNA viruses and regulate the viral replication [9]. The regulatory functions of miRNAs are further regulated by circRNAs and thus form a tight layer of competing endogenous RNA network (ceRNA) to regulate various signaling pathways under different physiological conditions. The role of circRNAs in regulating innate antiviral immune response against viral infection is not well-studied. Most circRNA studies are related to DNA viruses, and no previous studies have explored the mechanistic role of circRNAs in DENV infection [10, 11].

In this study, after the comprehensive computational analysis of the high-depth clinical RNA-seq dataset obtained from Indian patients, circEPSTI1 was identified to be highly expressed and upregulated in the PBMCs of the Dengue patients. The upregulation was validated in the plasma of Indian Dengue patients recruited for this study. Furthermore, the characterization of circEPSTI1 revealed that its upregulation is interferon-dependent, and the knockdown and overexpression of circEPSTI1 suppress and enhance DENV replication, respectively. Mechanistically, circEPSTI1 exerts its pro-viral role by sponging miR-942-5p, a miRNA that suppresses DENV replication by targeting the DENV genomic RNA at the NS1, NS3, and NS5 loci, as well as the host AKT1 signaling pathway. Using high-throughput RNA-sequencing, SERPINE1 expression was found to be dysregulated in the circEPSTI1 knockdown samples, and the significant gene signature resembles that of the AKT1 knockout mouse. circEPSTI1 sponging of miR-942-5p relieves its suppression on SERPINE-1, which in turn, activates the AKT1 signaling pathway. AKT1 signaling is crucial for the cellular survival that DENV hijacks for replication, thus facilitating the viral propagation. Thus, circEPSTI1 functions as a competing endogenous RNA that modulates the host antiviral response by regulating the activity of miR-942-5p during DENV infection.

## Results

### circEPSTI1 induction is observed in the plasma of dengue patients and in different cell types in an interferon-dependent manner

To quantify the reliable circRNA expression in the clinical samples of dengue patients, public sequencing databases were mined. After screening for datasets with non-polyA-enriched total RNA libraries and high sequencing depth, only the dataset GSE94892 remained. The read distribution analysis of the human mapped reads, unmapped reads, and the circRNA reads in every sample revealed the accurate and reliable circRNA detection (**Figure 1A)**. Principal Component Analysis (PCA) revealed a clear segregation between the control and DENV-infected groups based on circRNA expression **(Supplementary Figure 1A)**, indicating distinct transcriptomic alterations during infection. Differential expression analysis revealed 180 dysregulated circRNAs (142 upregulated, 36 downregulated; padj < 0.05, |log FC| > 1) in the dengue-infected patient group in comparison to the control group **(Figure 1B)**. Among the top significant circRNAs, circEPSTI1 (hsa_circEPSTI1_001 – CircBank, hsa_circ_0000479 - CircBase) was consistently upregulated and highly expressed across samples, and hence selected for further validation **(Figure 1C, Supplementary Figure 1B–C)**.

**Figure 1:**
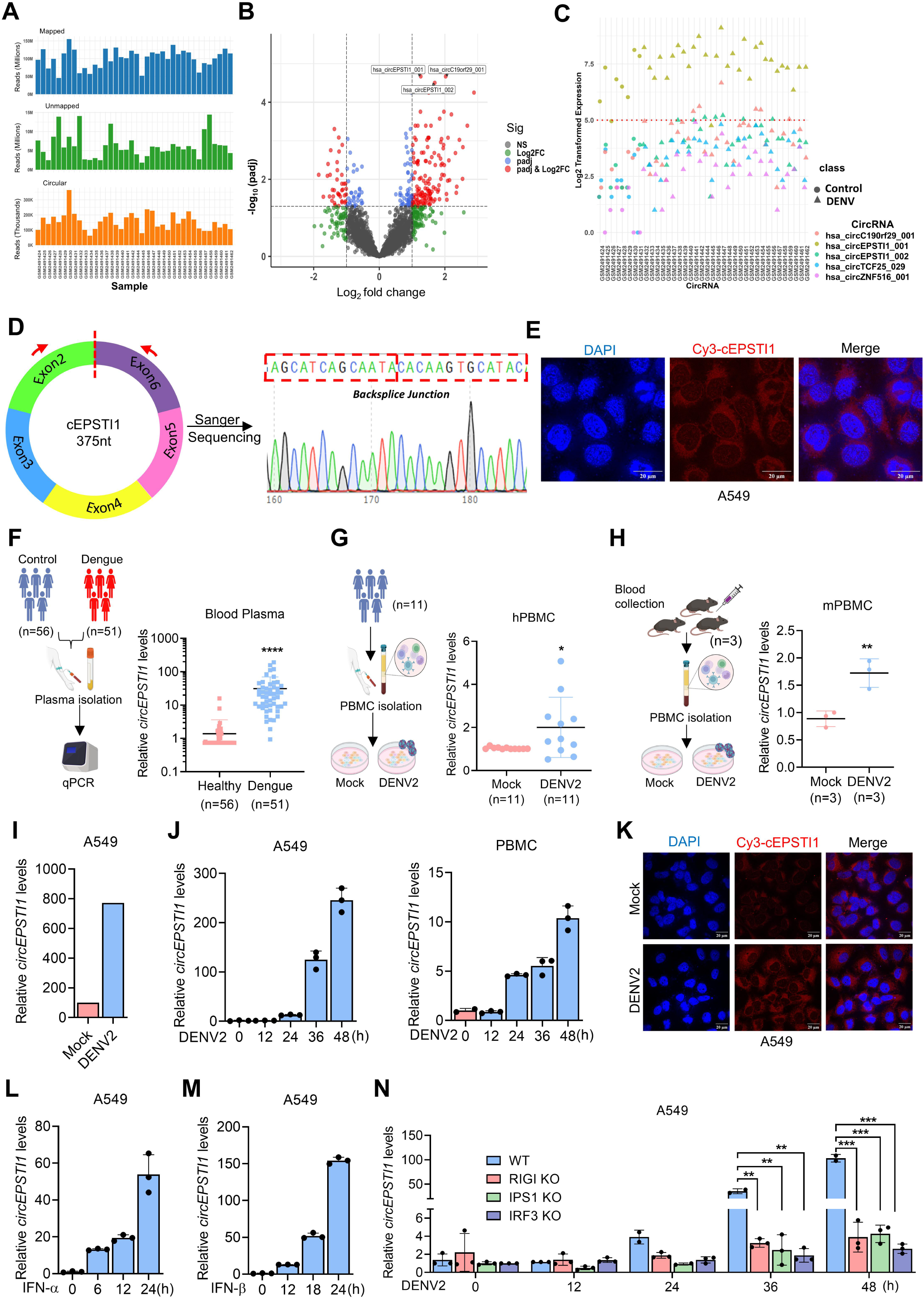
circRNA Profiling identifies circEPSTI1 to be induced upon DENV infection in an interferon-1 dependent manner. (A) Distribution of sequencing reads mapped to the human genome: mapped reads (blue), unmapped reads (green), both in millions, and circular reads (orange), in thousands across samples from dataset GSE94892, supporting high-depth circRNA profiling in DENV-infected human PBMCs. (B) Volcano plot showing differentially expressed circRNAs between control and DENV-infected groups. Red points indicate statistically significant circRNAs based on log fold change and adjusted p-values. The top three circRNAs are annotated with arrows. (C) Dot plot depicting normalized expression levels of the top five circRNAs. The red dotted line is the threshold with the normalized value of 5, above which the circRNAs are expressed at a higher level. (D) Sanger sequencing of the amplified bands using specific divergent primers spanning the back-splice junction. (E) FISH analysis using Cy3-labeled probe against circEPSTI1 of DENV2-infected A549 cells for localization of circEPSTI1. (F) Schematic of clinical recruitment of DENV patients’ plasma from India. circEPSTI1 levels were quantified by qPCR in healthy controls (n=56) and DENV (n=51) patients. (G) Human PBMCs (n=11) of healthy volunteers were infected with DENV (MOI=2) for 36h ex vivo, and circEPSTI1 levels were quantified by qPCR. (H) Mouse PBMCs (n=3) were infected with DENV (MOI=4) for 36h ex vivo, and circEPSTI1 levels were quantified by qPCR. (I) The copy number of circEPSTI1 was quantified using digital-PCR in the DENV-infected (MOI=0.5) and mock-infected A549 cells at 36h. (J) A549 and primary cells, PBMCs, were infected with DENV, and the circEPSTI1 level was quantified at 12, 24, 36, and 48h post-infection by qPCR. (K) Mock and DENV2 infected A549 cells were fixed and hybridized with Cy3-labelled circEPSTI1 probe and subjected to confocal microscopy. A549 cells were treated with (0.5ug/ml) recombinant IFN-α (L) and (0.5ug/ml) recombinant IFN-β (M), and the level of circEPSTI1 was quantified at different time points by qPCR. (N) DENV infection (MOI=0.5) was performed in wild-type (WT), RIG-I (Sensor) Knock-out (KO), IPS-1 (Adaptor) KO, or IRF-3 (transcription factor) KO A549 cells at 12, 24, 36, and 48 h, and the expression of circEPSTI1 was quantified by qPCR. Data are mean ± SEMs from triplicate samples of a single experiment and are representative of results from three independent experiments.

circEPSTI1 is derived through the back-splicing between the exon 2 and exon 6 of EPSTI1 pre-mRNA, leading to the 375 bp circular transcript (**Supplementary Figure 2A)**. Divergent primers spanning the back-splice junction specifically amplified the circEPSTI1, confirmed by Sanger sequencing **(Figure 1D)**. Divergent primers amplified the circEPSTI1 from random hexamer-derived cDNA, but not genomic DNA (gDNA) or oligo-dT cDNA, indicative of circRNA property, while convergent primers targeting linear EPSTI1 amplified in genomic DNA and oligo-dT cDNA of A549 and PBMCs **(Supplementary Figure 2B)**. Actinomycin D and RNaseR treatment also did not affect circEPSTI1 expression, demonstrating its stability, while linear EPSTI1 reduction was observed in A549 and PBMCs **(Supplementary Figure 2C, 2D)**. Confocal microscopy imaging performed through Fluorescence in situ Hybridization (FISH) utilizing a Cy3-labelled probe targeting circEPSTI1 identified the predominant localization in the cytoplasm **(Figure 1E)**. This finding was further supported by the subcellular fractionation assay in A549 and PBMCs **(Supplementary Figure 2E)**.

To validate the findings, plasma from dengue patients and healthy controls was analyzed by RT-PCR **(Supplementary Table S1)**. A significant induction of circEPSTI1 was observed in dengue patients compared to healthy controls **(Figure 1F)**. circEPSTI1 upregulation was also observed in the human PBMCs infected ex vivo with DENV2 **(Figure 1G) (Supplementary Figure 3A)**. Given its conservation in mice **(Supplementary Figure 3B)**, ex vivo infection of mouse PBMCs confirmed similar circEPSTI1 induction, indicating a conserved response across species **(Figure 1H) (Supplementary Figure 3C)**. The copy number of circEPSTI1 was found to be >750 copies/µl in DENV-infected A549 cells compared to the uninfected cells **(Figure 1I)**. Time-course analysis of DENV infection in multiple cell types, including A549, PBMC **(Figure 1J)**, HepG2, and MRC5 **(Supplementary Figure 3D & 3E),** was performed to understand the dynamics of circEPSTI1 induction. Progressive induction was observed at 24, 36, and 48 hours post-infection in all cell types. The cellular localization of circEPSTI1 was imaged upon DENV infection by confocal microscopy, and significant enrichment of circEPSTI1 was observed in the cytoplasm of DENV-infected cells **(Fig. 1K)**. Upon DENV infection, the primary innate antiviral response is initiated by RLR signaling, which induces type I interferons and inflammatory cytokines. Upon treatment with type I interferons, IFN-α and IFN-β, circEPSTI1 expression was induced, suggesting a role for interferon-mediated induction **(Figure 1L, 1M)**. To explore the involvement of type-I interferon signaling for circEPSTI1 induction and whether it is RLR-dependent, the innate immune RLR sensor, RIG-I, RLR signaling adaptor IPS-1, and the downstream interferon-inducing transcription factor, IRF3, were knocked out using the CRISPR-Cas9 technique in A549 cells. circEPSTI1 was induced in WT cells but not in the cells deficient in RIG-I, IPS-1, and IRF3, suggesting the interferon-dependent, RLR-dependent induction of circEPSTI1 **(Figure 1N)**. Taken together, circEPSTI1 induction upon DENV infection is observed to be interferon-dependent and is predominantly expressed in the cytosolic compartment.

### circEPSTI1 regulates DENV infection

The role of circEPSTI1 in DENV replication was explored using siRNA-mediated circEPSTI1 knockdown. siRNA targeting the back-splice junction of circEPSTI1 significantly reduced the expression of circEPSTI1 in PBMCs, A549, and MRC5 cells **(Figure 2A)**, while linear EPSTI1 expression remained unchanged **(Supplementary Figure 3F)**. Knockdown of circEPSTI1 significantly suppressed the viral load in PBMCs, A549, and MRC5 **(Figure 2B)** and in the culture supernatant of A549 **(Figure 2C)**. Further, DENV replication was suppressed upon circEPSTI1 knockdown as observed through western blot **(Figure 2D)**, flow cytometry **(Figure 2E)**, and confocal microscopy **(Figure 2F)** using dengue NS5-specific antibody. Collectively, our results with multiple techniques demonstrate that circEPSTI1 knockdown suppresses DENV replication. To strengthen the findings, circEPSTI1 overexpression was performed by cloning circEPSTI1 into the mc2mNeonV2 vector, and the overexpression was verified by RT-PCR, which shows >1400-fold increase in circEPSTI1 **(Figure 2G)**. DENV viral load was increased to a higher extent upon circEPSTI1 overexpression in A549 cells **(Figure 2H)** and in culture supernatant **(Figure 2I).** The finding was further validated through western blot **(Figure 2J),** flow cytometry **(Figure 2K)**, and confocal microscopy **(Figure 2L)**. Altogether, the DENV viral load increased upon circEPSTI1 overexpression, as demonstrated by various techniques.

**Figure 2:**
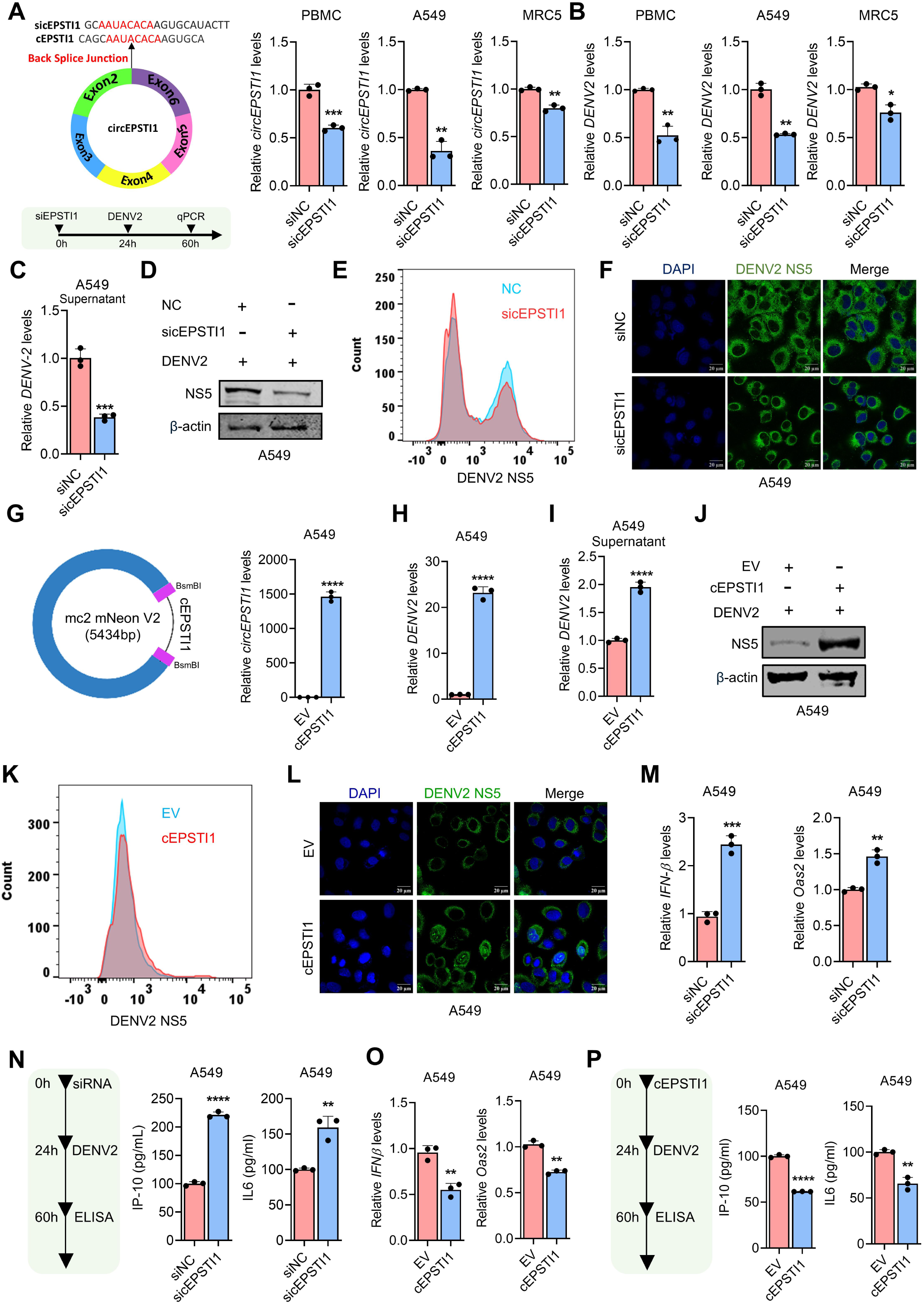
DENV establishes infection through circEPSTI1. (A) siRNA-mediated knockdown of circEPSTI1 was performed by designing the siRNA against the back-splice junction, as shown in the schematic. PBMCs, A549, and MRC5 cells were transfected with 25nM of circEPSTI1 siRNA for 24h, and subsequently, the cells were infected with DENV for 36h. Knockdown efficiency of circEPSTI1 and linear EPSTI1 was quantified by qPCR. (B) Intracellular viral load was quantified by qPCR. (C) Viral load in the culture supernatant was quantified upon circEPSTI1 knockdown by qPCR in A549 cells. A549 cells transfected with circEPSTI1 siRNA were infected with DENV (MOI=0.5) for 36h. DENV viral load at the protein level (NS5) upon circEPSTI1 knockdown was quantified using the (D) western blot, (E) FACS analysis, and (F) confocal microscopy using DENV NS5-specific antibody. Nuclei were stained with DAPI. (G) circEPSTI1 was cloned into the Circminivector mneon v2 vector, as shown in the schematic. A549 cells were transfected with the overexpression vector and infected with DENV (MOI=0.5) for 36h, and overexpression was tested by q-PCR. A549 cells transfected with the circEPSTI1 vector were infected with DENV (MOI=0.5) for 36h. Viral load in (H) A549 cells and (I) culture supernatant was quantified by qPCR. DENV NS5 protein level in circEPSTI1 was quantified through (J) western blot analysis, (K) FACS analysis, and (L) confocal microscopy using DENV NS5 antibody. Nuclei were stained with DAPI. Upon circEPSTI1 knockdown, the levels of (M) IFNβ and OAS2 were quantified by qPCR, and the levels of (N) interferon-induced protein IP-10 and inflammatory cytokine IL-6 were quantified using ELISA. upon (M) circEPSTI1 knockdown and (N) circEPSTI1 overexpression in A549 cells. Upon circEPSTI1 overexpression, the levels of (O) IFNβ and OAS2 were quantified by qPCR, and the levels of (P) interferon-induced protein IP-10 and inflammatory cytokine IL-6 were quantified using ELISA in A549 cells. PBMC, A549, and MRC5 cell lines were infected by DENV2 at a MOI of 2, 0.5, and 1, respectively. Data are mean ± SEMs from triplicate samples of a single experiment and are representative of results from three independent experiments. ∗∗∗∗, p < 0.0001, ∗∗∗, p < 0.001, ∗∗, p < 0.01, and ∗, p < 0.05, by two-tailed unpaired t test (B). ns, nonsignificant.

Next, the effect of circEPSTI1 on innate antiviral responses was investigated. circEPSTI1 knockdown significantly increased the expression of IFN-β and interferon-induced gene, OAS2 **(Figure 2M),** and enhanced the secretion of IFN-inducible protein, IP-10, and inflammatory cytokine IL-6, in culture supernatant **(Figure 2N).** Conversely, circEPSTI1 overexpression suppressed the expression of IFN-β and OAS2 (Figure 2O), as well as the secretion of IP-10 and IL-6 **(Figure 2P)**. Overall, knockdown of circEPSTI1 reduces the viral load and increases IFNβ and inflammatory cytokines, whereas overexpression of circEPSTI1 increases the viral load and decreases the level of IFNβ and inflammatory cytokines, suggesting the pro-viral role of circEPSTI1 facilitating DENV replication.

### circEPSTI1 sponges miR-942-5p during DENV infection

Circular RNAs exert their regulatory functions mainly by sponging miRNAs to regulate the target genes. To identify the miRNAs that can bind to the circEPSTI1, three algorithms-miRanda, PITA, and TargetScan were utilized. miR-942-5p was the top miRNA candidate among all the tools, with the lowest free energy, highest miRanda score, and strongest complementarity **(Figure 3A)**. To validate the predictions, the circRNA sequence was cloned downstream of the luciferase gene under the CMV promoter. Upon overexpression of miR-942-5p, the luciferase activity was reduced significantly, whereas endogenous inhibition of miR-942-5p abolished this effect, suggesting the suppression is not due to non-specific binding **(Figure 3B)**. To further validate the findings, the predicted binding sites of miR-942 in the circ-EPSTI1 sequence were mutated, and this mutated construct failed to reduce the luciferase activity upon miR-942-5p overexpression, suggesting that miR-942 binds specifically to circEPSTI1 with its seed sequence at the predicted sites **(Figure 3C)**. The miRNA targets the transcript through a multiprotein complex known as RNA-induced silencing complex (RISC), and Argounate-2 protein (AGO2) is one of the essential components of RISC for miRNA-mediated silencing (**Supplementary** Fig. 4A). To explore the involvement of AGO2 in circEPSTI1 and hsa-miR-942-5p interaction, the AGO2 pulldown assay was performed using a FLAG-specific antibody (**Supplementary** Fig. 4B). The cells transfected with miR-942 showed > 4-fold enhancement of circEPSTI1 levels in the miR-942-5p transfected group compared to the NC-transfected group **(Figure 3D)**. circEPSTI1 was pulled down using a biotin probe, and miR-942-5p levels were enriched in the pull-down fraction in the DENV-infected cells, suggesting the direct interaction of circEPSTI1 and hsa-miR-942-5p **(Figure 3E) (Supplementary** Fig. 4C). Colocalization of miR-942-5p and circEPSTI1 was observed by confocal microscopy in DENV-infected A549 cells **(Figure 3F)**. miR-942-5p levels were significantly upregulated upon circEPSTI1 knockdown in PBMCs and A549 cells compared to the negative control transfected cells **(Figure 3G),** suggesting the possible inverse regulatory relationship between circEPSTI1 and hsa-miR-942-5p. This implies that the pro-viral activity of circEPSTI1 is mediated through the regulation of hsa-miR-942-5p expression. Furthermore, miR-942-5p expression was suppressed at different time points following DENV infection in PBMC, A549, MRC5, and HEPG2 cell lines, and was negatively correlated with the induction of circEPSTI1 expression **(Figure 3H)**. The suppression of miR-942-5p levels in dengue-infected cells was observed through confocal microscopy **(Figure 3I).** Overall, these findings suggest that circEPSTI1 sponges miR-942-5p upon DENV infection, and their interaction may be responsible for the pro-viral activity of circEPSTI1.

**Figure 3:**
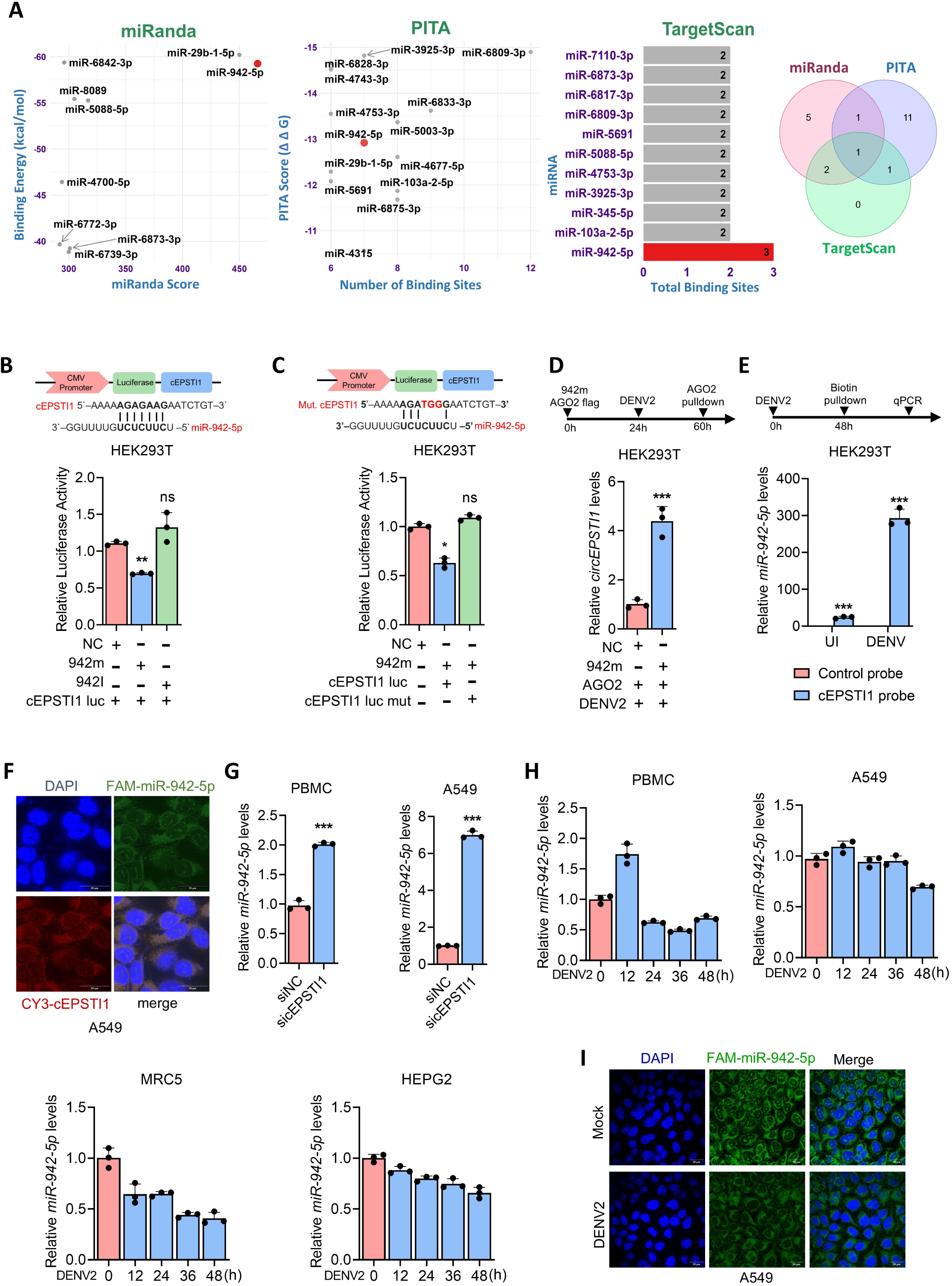
CircEPSTI1 sponges hsa-miR-942-5p. (A) Three RNA interaction prediction algorithms, miRanda, PITA, and TargetScan, were employed to predict the interaction of miRNAs with circEPSTI1 with a stringent cutoff: miRanda - Binding energy cutoff −40 kcal/mol and minimum miRanda Score of 300; PITA - PITA score of minimum −10 and number of binding sites to be minimum 6; TargetScan - more than one binding site. (B & C) circEPSTI1 wild type (WT) and mutant (M) sequences were cloned downstream of luciferase. HEK293T cells were transfected with indicated, 100ng of circEPSTI1-luc (WT) or mut circEPSTI1 (M), 25nM miR-942-5p mimic or 25nM miR-942-5p inhibitor, and 30ng pRL-TK vector. Luciferase assay was performed after 48h. (D) HEK293T cells were transfected with 2µg of plasmid encoding Flag-Ago2 and 25nM mir-942 and were infected with DENV virus (MOI=2) for 36h. RNA immunoprecipitation was done using FLAG-M2 beads, and the enrichment of miR-942 was quantified by qPCR. (E) HEK293T cells were infected with DENV for 36 hrs. A biotin-labeled circEPSTI1 probe was used to pull down the RNA fractions binding to the circEPSTI1, and the level of miR-942 was quantified by qPCR. (F) A Cy-3 labelled probe against circEPSTI1 and a FAM-labelled probe against miR-942-5p were used to visualize the colocalization of both by FISH analysis in A549 cells. (G) Upon knockdown of circEPSTI1, miR-942-5p expression was quantified in PBMC and A549 cells by qPCR. (H) miR-942-5p levels upon DENV infection at different time points were quantified in PBMC (MOI =2) and A549 (MOI=0.5), MRC5(MOI=0.5), and HEPG2(MOI=0.5) by qPCR. (I) Mock and DENV2-infected A549 cells were fixed and hybridized with FAM-labelled miR-942 probe and subjected to FISH analysis. Data are mean ± SEMs from triplicate samples of a single experiment and are representative of results from three independent experiments. ∗∗∗∗, p < 0.0001, ∗∗∗p < 0.001 ∗∗, p < 0.01, and ∗p < 0.05, by two-tailed unpaired t-test (B). ns, nonsignificant.

### hsa-miR-942-5p inhibits DENV replication by targeting the viral genome

MicroRNAs bind to the genome of RNA viruses such as Influenza and regulate their replication [9, 12]. The miRanda tool was used to determine whether miR-942-5p can bind to the DENV genomic RNA across all four serotypes **(Figure 4A)**. Based on the conservation of the site across more than one DENV serotype and the free energy, eight sites in the DENV genome were identified. NS5 bears 4 sites, NS1 bears 3 sites, NS3 bears 2 sites, and E bears a single site **(Figure 4B)**. To determine the involvement of AGO2-mediated miR-942-5p and DENV viral RNA interaction, AGO2 pulldown was performed, and DENV RNA was enriched in miR-942-5p transfected group compared to the NC-transfected group in A549 **(Figure 4C)** and HEK293T cells (**Supplementary** Fig. 5A). To validate the prediction, NS5, NS3, and NS1 were cloned downstream of the luciferase, and the luciferase activity upon miR-942-5p overexpression was quantified. NS1 luciferase activity was suppressed the most, followed by NS3 and NS5. When NS2B with no predicted binding site was cloned downstream of luciferase, luciferase activity suppression was not observed, suggesting the predictions are not random and non-specific **(Figure 4D, 4E, 4F & 4G).** miR-942-5p overexpression **(Supplementary Figure 5B)** and suppression **(Supplementary Figure 5C)** in A549 cells were performed using synthetic mimics and inhibitors, respectively. While miR-942-5p overexpression suppressed the DENV, the same effect was not observed upon the transfection of miR-942-5p inhibitor in A549 and PBMCs **(Figure 4H)** and in the A549 supernatant **(Supplementary** Fig. 5D). The level of NS1 and NS5 at the protein level was evaluated through the western blot. DENV protein levels were reduced upon miR-942-5p overexpression, suggesting the anti-viral role of miR-942-5p upon DENV infection **(Figure 4I)**. Further, the reduction of DENV NS1 and NS5 was validated by FACS **(Figure 4J)** and immunofluorescence through confocal microscopy **(Figure 4K)** using NS1 and NS5-specific antibodies. Further, the levels of IFNβ interferon-inducible protein, IP-10, and inflammation-associated cytokines such as IL-6 were quantified through ELISA. Upon miR-942-5p overexpression, IL-6 and IP-10 levels were significantly induced **(Figure 4L)**. Overall, miR-942-5p suppresses DENV infection by primarily targeting the DENV NS1, NS3, and NS5 to modulate the host innate immune response. Notably, simultaneous inhibition of circEPSTI1 and miR-942-5p could not suppress the viral load, as observed in circEPSTI1 alone, suggesting the miR-942-5p-mediated viral suppression of circEPSTI1 **(Supplementary** Fig. 5E).

**Figure 4:**
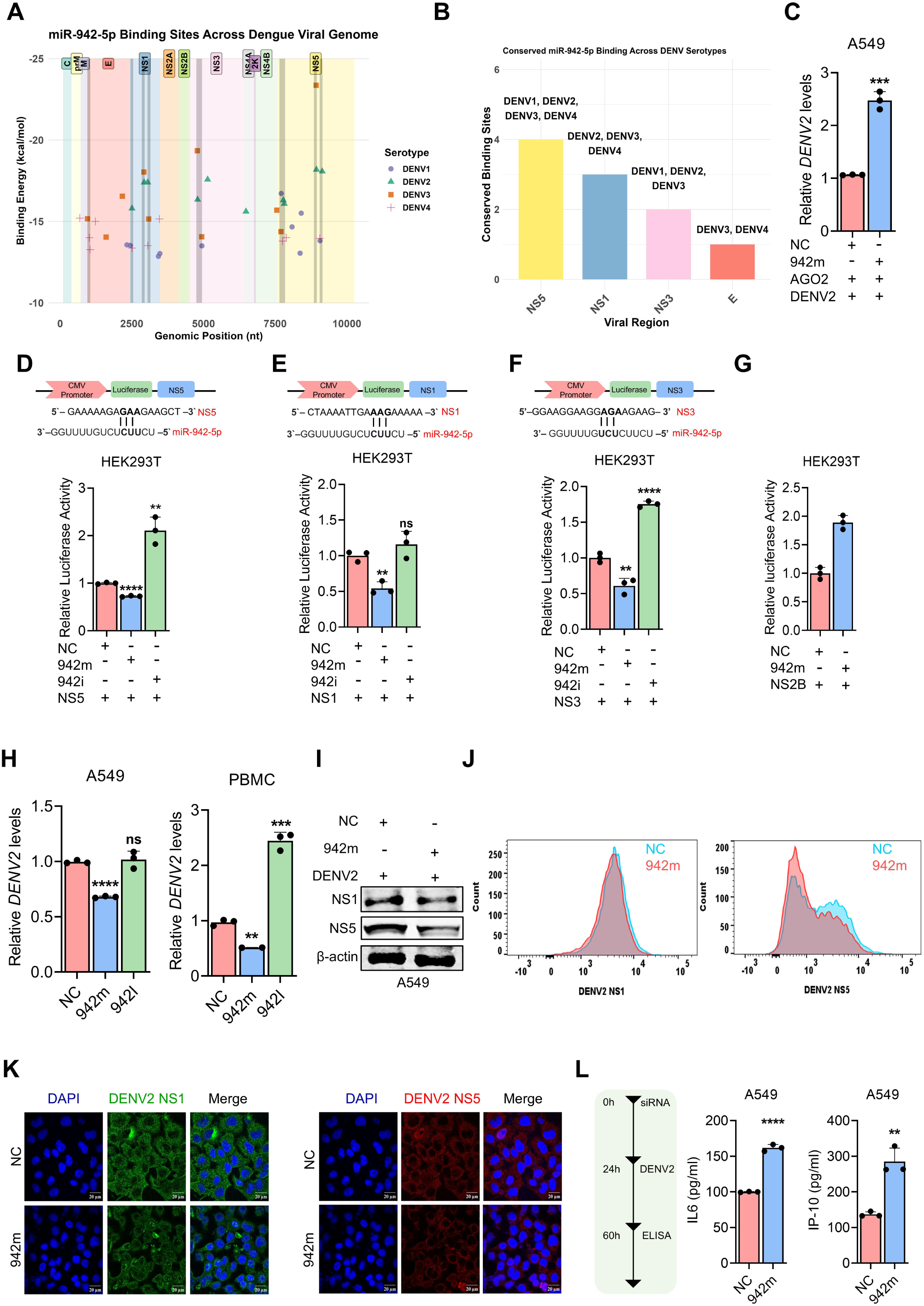
miR-942-5p targets DENV genomic RNA. (A) Predicted binding sites of miR-942-5p across the genomes of four different DENV serotypes. (B) The number of conserved binding sites of miR-942-5p across the DENV viral genome. (C) A549 cells were transfected with 2µg of FLAG-Ago2 plasmid and 50 nM of miR-942 mimic, followed by DENV infection for 36h. RNA immunoprecipitation was done using FLAG-M2 beads, and the enrichment of the virus was tested by qPCR. (D, E, F & G) NS5, NS1, NS3, and NS2B sequences were cloned downstream of the pmiR-luciferase vector. HEK293T cells were transfected with 100 ng of the luciferase constructs, 25nM of miR-942 mimic, 25nM of miR-942 inhibitor, and 30ng of PRL-Tk plasmid. Luciferase assay was performed after 48h. (H) A549 cells and PBMCs were transfected with 25nM of miR-942 mimic and 25nM of miR-942 inhibitor for 24h and infected with DENV for 36h. The viral load was quantified by qPCR. (I) DENV NS1 and NS5 protein level in miR-942 overexpressing A549 cells was quantified through western blot analysis. (J) FACS analysis was performed with the DENV NS1 and NS5-specific antibody upon miR-942 overexpression in A549 cells. (K) A549 cells transfected with miR-942 mimic were infected with DENV for 36h and were subjected to confocal microscopy using DENV NS1 and NS5 antibodies. Nuclei were stained with DAPI. (L) The levels of interferon-induced protein IP-10 and inflammatory cytokine IL-6 were quantified using ELISA upon miR-942 overexpression in A549 cells. Data are mean ± SEMs from triplicate samples of a single experiment and are representative of results from three independent experiments. ∗∗∗∗, p < 0.0001, ∗∗∗p < 0.001 ∗∗, p < 0.01, and ∗p < 0.05, by two-tailed unpaired t-test (B). ns, nonsignificant.

### circEPSTI1 sponges miR-942-5p to regulate the SERPINE1

To understand how circEPSTI1 affects host gene expression, high-throughput RNA-sequencing was performed upon knockdown of circEPSTI1. PCA plot showed a clear segregation between the Negative control (NC_DENV) and circEPSTI1 knockdown DENV-infected (sicircEPSTI1_DENV) group **(Figure 5A)**. The differentially expressed genes were visualized using a volcano plot **(Figure 5B)**. SERPINE1, MBOAT1, SORL1, and CBS were the common genes among the miR-942-5p targets, and the genes downregulated by circEPSTI1 knockdown **(Figure 5C)**. SERPINE1 was selected for further analysis due to the higher target score and significant dysregulation among the other three genes in the RNA-seq data analysis **(Figure 5D)** (**Supplementary** Fig. 6A). SERPINE1/PAI-1 was reported to play diverse roles in the regulation of platelet biogenesis and cell-death-associated pathways. SERPINE1 level was significantly increased in dengue patients compared to the healthy group **(Figure 5E)**. SERPINE1 expression was progressively induced over time post DENV infection at protein level **(Figure 5F)** and RNA level (**Supplementary** Fig. 6B) and IFNβ induction (**Supplementary** Fig. 6C). Reduction in the expression level of SERPINE1 in circEPSTI1 knockdown samples was validated at the RNA level using RT-PCR **(Figure 5G)** and at the protein level through western blot **(Figure 5H),** and confocal microscopy using SERPINE-1 specific antibody **(Figure 5I)**.

**Figure 5:**
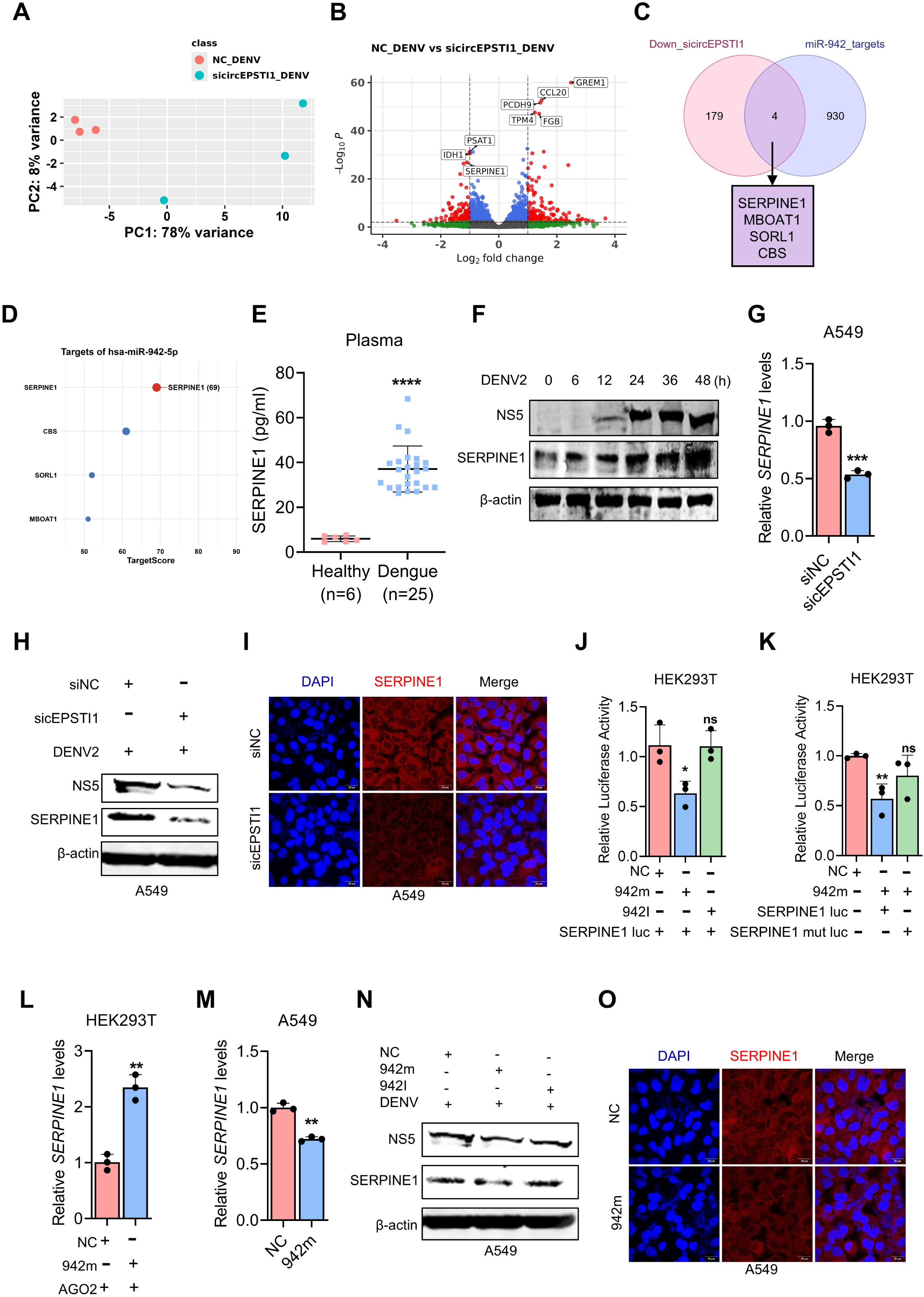
circESPTI1 inhibits miR-942-5p binding to host SERPINE1. (A) PCA plot showing the segregation of NC vs sicircEPSTI1 transfected DENV infected groups. (B) Volcano plot showing the differentially expressed genes between the Negative control transfected DENV-infected (NC_DENV) and the sicircEPSTI1 transfected DENV-infected (sicircEPSTI1_DENV) groups. (C) Venn diagram showing the common genes between circEPSTI1 knockdown and miR-942-5p targets. (D) Predicted Targetscore of the genes binding with miR-942-5p. (E) The levels of SERPINE1 were quantified in healthy and Dengue patients by ELISA. (F) A549 cells were infected with DENV at MOI=0.5 and the levels of SERPINE1 were checked at different time points by Western Blot Analysis. (G) A549 cells transfected with 25nM circEPSTI1 siRNA were infected with DENV (MOI=0.5) for 36h, and the levels of SERPINE1 were quantified by qPCR. (H) The levels of SERPINE1 at the protein level were quantified by Western blot analysis and (I) confocal microscopy upon circEPSTI1 knockdown in A549 cells. (J) HEK293T cells were transfected with 100ng of circEPSTI1-luc, 25nM miR-942-5p mimic or 25nM miR-942-5p inhibitor, and 30ng pRL-TK vector. Luciferase assay was performed after 48h. (K) HEK293T cells were transfected with 100ng of circEPSTI1-luc or mut circEPSTI1, 25nM miR-942-5p mimi, and 30ng pRL-TK vector. Luciferase assay was performed after 48h. (L) HEK293T cells were transfected with 2µg of FLAG-Ago2 plasmid and 50 nM of miR-942 mimic, followed by DENV infection for 36h. RNA immunoprecipitation was done using FLAG-M2 beads, and the levels of SERPINE1 were quantified by qPCR. (M) A549 cells transfected with 25nM of miR-942 mimic were infected with DENV (MOI=0.5) for 36h, and the levels of SERPINE1 were quantified by qPCR. (N) A549 cells were transfected with 25nM of miR-942 mimic for 24h and infected with DENV for 36h. SERPINE1 protein level was quantified through western blot analysis and (O) confocal microscopy.

To validate the binding between the miR-942-5p and SERPINE1, the 3’-UTR of SERPINE1 was cloned downstream of luciferase, and upon miR-942-5p overexpression, the luciferase activity was reduced, whereas upon miR-942-5p inhibitor transfection, no change was observed **(Figure 5J)**. When the miR-942-5p binding sites in the SERPINE1 3’-UTR were disturbed, there was no change observed, suggesting the specific binding of miR-942-5p with SERPINE1 UTR **(Figure 5K)**. Significant enrichment of SERPINE1 was observed in the miR-942-5p overexpressed AGO2-pulldown fraction **(Figure 5L)**. Moreover, SERPINE1 expression downregulation upon miR-942-5p overexpression was observed at the RNA level using RT-PCR **(Figure 5M)**, protein level using western blot **(Figure 5N)**, and the confocal microscopy using SERPINE1-specific antibody **(Figure 5O).**

Overall, circEPSTI1 and miR-942-5p regulate the expression of SERPINE1, and this axis is dysregulated in dengue infection.

### Tiplaxtinin targets the pro-viral circEPSTI1-miR-942-5p-SERPINE1 axis to suppress DENV replication

To investigate the downstream pathways regulated by the circEPSTI1-miR-942-5p-SERPINE1 axis, pathway-level analyses were performed. Interestingly, 214 out of 836 significantly downregulated genes upon circEPSTI1 knockdown were also found to be downregulated in the AKT-1 knockout mouse from the public dataset GSE39699, as determined by reverse signature analysis **(Figure 6A)**. miR-942-5p target gene enrichment analysis revealed that the PI3K-Akt signaling pathway is one of the top dysregulated pathways. Since circEPSTI1 knockdown exhibits a signature similar to AKT1 knockout and miR-942-5p targets enrich for the PI3K-AKT1 signaling pathway (Figure 6B), we speculated that SERPINE1 may also be involved in the same signaling axis. Moreover, SERPINE1 was previously reported in a study to be an activator of the PI3K-AKT1 signaling pathway. Upon overexpression of SERPINE1, p-AKT levels increased, suggesting activation of the pathway, as observed in a previous study **(Figure 6C)**. AKT1 signaling plays a role as a pro-viral or anti-viral signaling axis during different virus infections. The levels of p-AKT1 increased progressively at different time points following DENV infection, indicating the activation of this pathway **(Figure 6D)**. Upon siRNA-mediated knockdown of AKT1, dengue viral load was suppressed at the RNA level **(Figure 6E)**, dengue NS5 protein level by western blot **(Figure 6F)**, FACS analysis **(Figure 6G)**, and confocal microscopy **(Figure 6H)**. Knockdown of AKT1 increased the expression of IFN-β and OAS2 (**Supplementary Figure 7A)**. The levels of cytokine IL-6 and IP-10, quantified by ELISA, increased in the supernatant **(Figure 6I)**.

**Figure 6:**
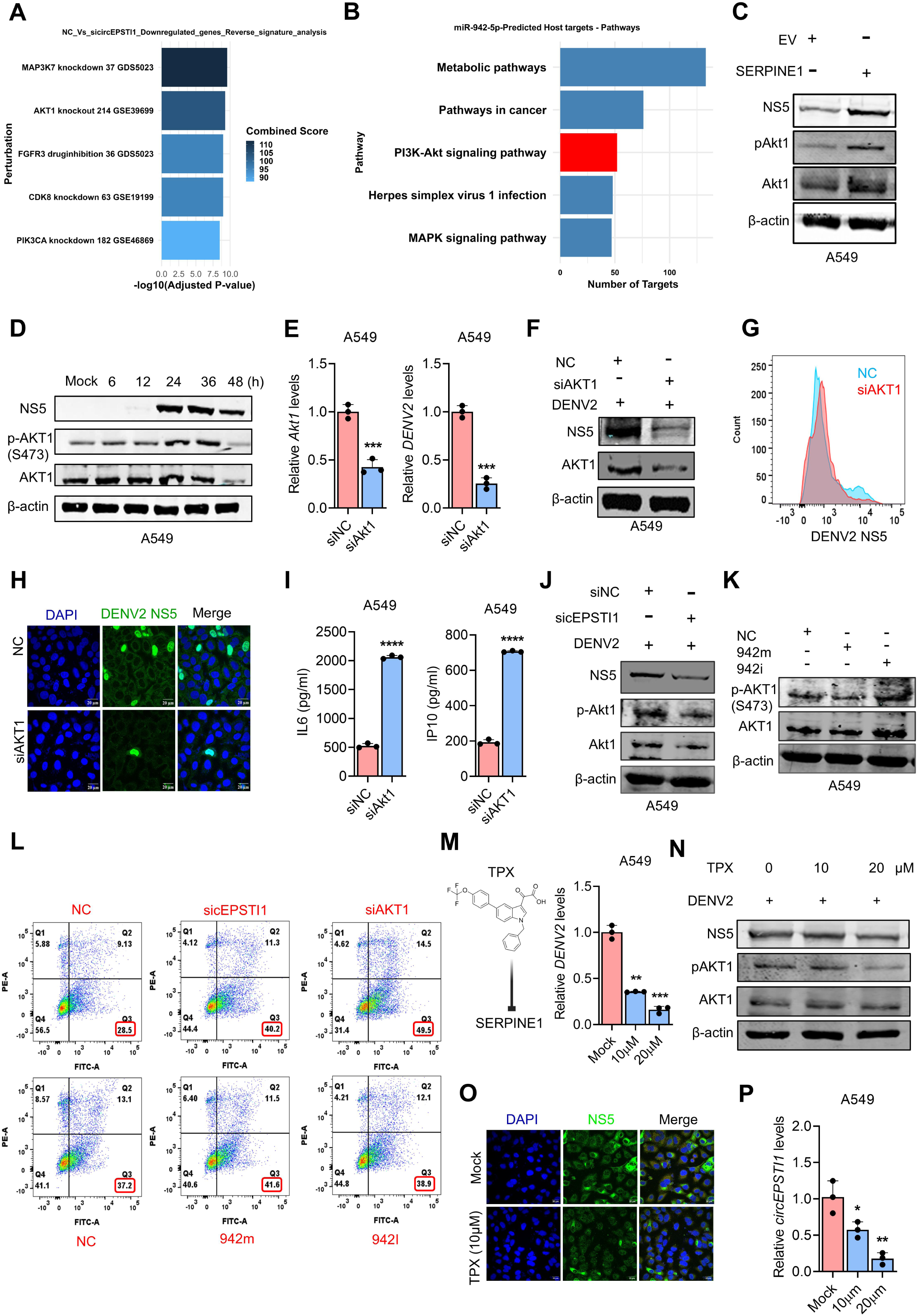
Tiplaxtinin reduces circEPSTI1 and dengue viral load through the suppression of SERPINE1-mediated AKT1 signaling. (A) Bar plot showing the top signatures resembling the differentially expressed genes upon circEPSTI1 knockdown. (B) The miRabel tool was utilized to perform the pathway enrichment analysis of miR-942-5p targets. (C) A549 cells transfected with 2μg of SERPINE1 plasmid were infected with DENV (MOI=0.5) for 36h, and the levels of AKT1 and p-AKT1 protein were quantified by western blot. (D) A549 cells were infected with DENV at an MOI of 0.5. Levels of NS5, AKT1, and p-AKT1 were quantified in DENV2-infected A549 cell line at different time points by Western blot analysis. (E) A549 cells were transfected with 25nM of AKT1 siRNA for 24h and infected with DENV for 36h. The knockdown efficiency and viral load were quantified by qPCR. (F) DENV NS5 protein level in siAKT1-transfected A549 cells was quantified through western blot analysis (G), FACS, and (H) confocal analysis was performed with the DENV NS5-specific antibody upon knockdown of AKT1 in A549 cells (I). The levels of interferon-induced protein IP-10 and inflammatory cytokine IL-6 were quantified using ELISA upon SERPINE1 overexpression in A549 cells. (J) A549 cells transfected with 25nM circEPSTI1 siRNA were infected with DENV (MOI=0.5) for 36h, and the levels of Akt1 and p-Akt1 protein were quantified by western blot. (K) A549 cells were transfected with 25nM miR-942 mimic and miR-942 inhibitor, and cells were infected with DENV for 36h. The levels of NS5, p-AKT1, and AKT1 were quantified by western blot analysis. (L) A549 cells transfected with 25nM circEPSTI1, AKT1 siRNA, and 25nM of 942 mimic and inhibitor were infected with DENV (MOI=0.5) for 36h. Cells were stained with Propidium Iodide (PI) alone and PI + annexin V/, and apoptosis was determined by using flow cytometric analysis. (M) A549 cells were pre-treated with 10 and 20 μM of Tiplaxtinin, a SERPINE1 inhibitor, for 2 hours, followed by infection with DENV (MOI=0.5) for 36h. The levels of DENV2 were quantified by qpCR. (N) The levels of DENV NS5, Akt1and pAKT1 protein levels were quantified by western blot in Tiplaxtinin-treated A549 DENV2-infected cells. (O) The viral load upon Tiplaxtinin treatment in A549 cells was quantified by confocal analysis. (P)The levels of circEPSTI1 in Tiplaxtinin-treated A549 cells were quantified by qPCR. Data are mean ± SEMs from triplicate samples of a single experiment and are representative of results from three independent experiments. ∗∗∗∗, p < 0.0001, ∗∗∗, p < 0.001, ∗∗, p < 0.01, and ∗, p < 0.05, by two-tailed unpaired t test (B). ns, nonsignificant.

To validate the findings from the reverse signature analysis of our RNA-seq data, which showed that circEPSTI1 knockdown resembles the signature of AKT1 knockout, siRNA-mediated knockdown of circEPSTI1 was performed, and p-AKT1 levels were quantified. circEPSTI1 knockdown decreases the levels of p-AKT1, suggesting the dampening of AKT1 signaling leading to reduced viral load **(Figure 6J)**. Similarly, miR-942-5p overexpression reduced the p-AKT1 activation. Conversely, sequestering endogenous miR-942-5p enhanced p-AKT1 activation **(Figure 6K)**. Overall, we can conclude that the circEPSTI1-miR-942-5p-SERPINE1 axis modulates the AKT1 signaling activation to regulate dengue infection.

As PI3K-AKT1 signaling is crucial for cellular survival, we hypothesized that DENV may utilize this pathway to enhance the viral load, and the regulation of this pathway is mediated by miR-942-5p. To test this hypothesis, siRNAs targeting circEPSTI1 and AKT1, as well as miR-942 mimics and inhibitors, were transfected. Following DENV infection, the cells were stained with Propidium Iodide (PI) alone and PI + annexin V, and apoptosis was determined using flow cytometric analysis. Knockdown of circEPSTI1 and AKT1 and overexpression of miR-942-5p induced apoptosis, suggesting the regulatory axis of circEPSTI1-miR-942-5p-AKT1 signaling in the regulation of DENV survival in infected cells (**Fig**. **6L**).

To identify the molecules that can target this axis and inhibit DENV replication, we first targeted p-AKT1 activation through the use of the AKT1 inhibitor wortmannin. Although Wortmannin treatment suppressed dengue viral load (**Supplementary Figure 7B)**, expression of circEPSTI1 did not change (**Supplementary Figure 7C)**, suggesting there are complex mechanisms through which AKT1 signaling facilitates DENV replication. Next, SERPINE1 inhibitor, Tiplaxtinin (TPX), treated cells were infected with DENV, and the effect was observed. TPX treatment reduced the DENV load at both the RNA level in A549 cells **(Figure 6M)** and PBMCs (**Supplementary Figure 7D)** and protein level through western blot **(Figure 6N)** and confocal microscopy **(Figure 6O)**. Interestingly, circEPSTI1 levels were also suppressed in a dose-dependent manner, suggesting the inhibitory effect of TPX over the circEPSTI1-miR-942-5p-SERPINE1 axis **(Figure 6P)** (**Supplementary Figure 7E)**. However, the inhibition of linear EPSTI1 was not observed (**Supplementary Figure 7F)**.

Overall, the induction of circEPSTI1 upon DENV infection enhances the viral load, resulting from the activation of AKT1 signaling through the regulation of miR-942-5p and SERPINE1. Interestingly, suppression of circEPSTI1 by Tiplaxtinin results in a reduction in viral load, indicating high potential for use as a dengue antiviral therapy by targeting this pathway **(Figure 7)**. These findings may also apply to other viral infections that exploit AKT1 signaling for replication.

**Figure 7:**
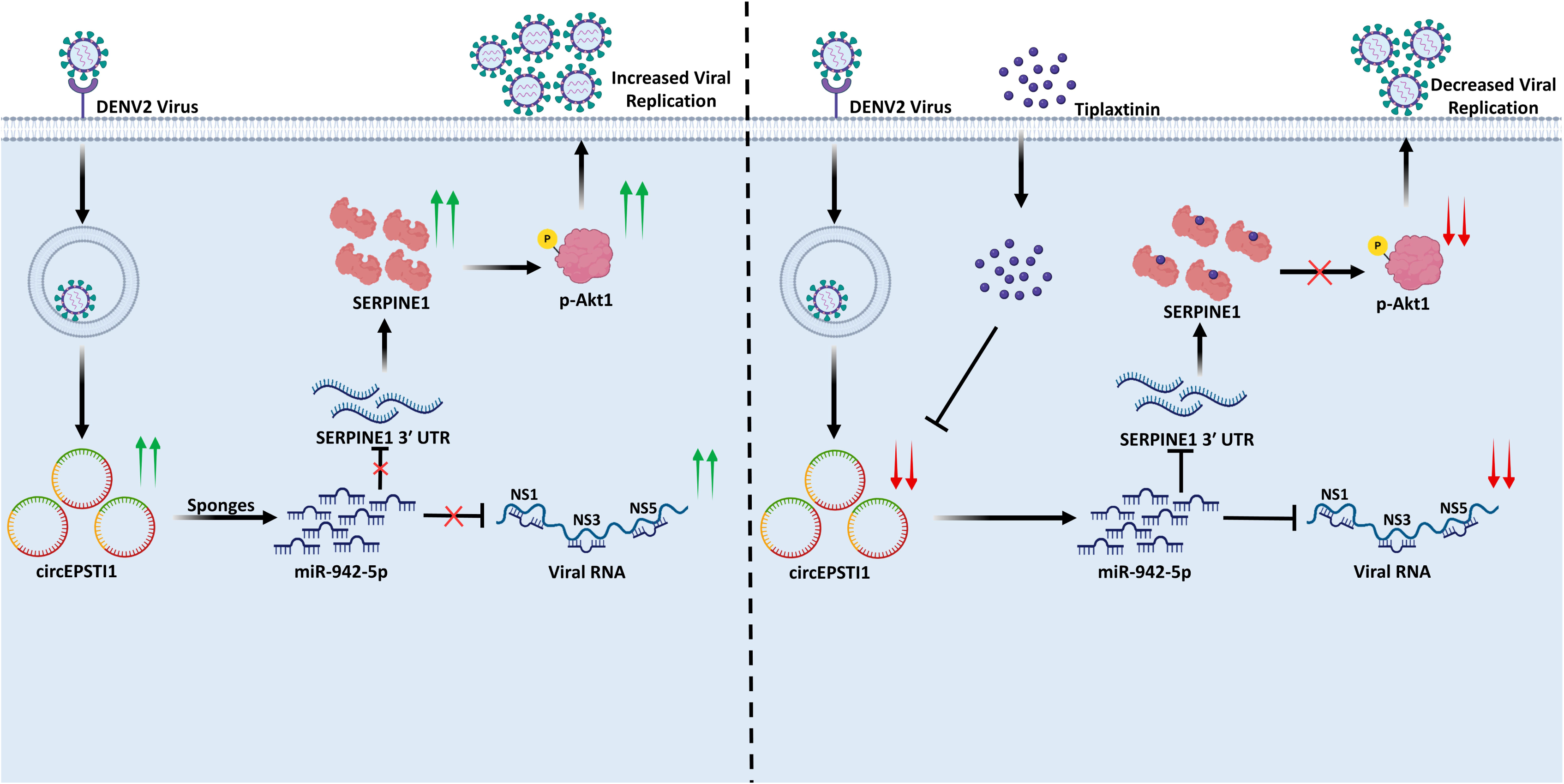
Schematic of the circEPSTI1 regulation of miR-942-5p-SERPINE1-AKT1 signaling to regulate DENV infection. Schematic representation of the mechanism by which circEPSTI1 sponges the miR-942-5p to relieve its suppression on SERPINE1 mRNA, leading to the activation of AKT1-signaling and increased viral replication. Meanwhile, inhibition of SERPINE1 by Tiplaxtinin reduces the expression of circEPSTI1 that leads to reduced activation of AKT1 signaling and suppressed dengue virus replication.

## Discussion

In this study, we provide the first evidence that circEPSTI1 is significantly induced and a host-derived proviral regulator of dengue infection. Upon DENV infection, the role of the circRNAs has not been previously studied. The only existing study identified two circRNAs to be dysregulated between pretreatment and post-treatment in whole blood samples of non-severe dengue patients and did not explore the functional role of any of the identified circRNAs [13]. circEPSTI1 was found to be upregulated in the plasma of severe and non-severe dengue patients compared to the healthy controls (**Fig. 1F**), and the induction was conserved in the mice PBMCs and across diverse human cell lines infected with DENV. Hence, circEPSTI1 is a highly promising diagnostic biomarker for dengue infection. The induction was RLR-signaling dependent, as the knockout of RIG-I, IPS-1, and IRF3 abolished the induction during DENV infection. Interestingly, unlike the classical interferon stimulatory genes, circEPSTI1 promotes viral replication.

Mechanistically, circRNAs often function by binding RNA-binding proteins (RBPs) or sponging miRNAs [3, 8]. Bioinformatic predictions, coupled with luciferase assays and AGO2 pull-down experiments, identified miR-942-5p as the primary miRNA target of circEPSTI1. The cytoplasmic localization of circEPSTI1 supports this sponging function. miR-942-5p has previously been implicated in antiviral responses during HCV infection [14], but its role in dengue was unknown. We demonstrate that miR-942-5p directly binds to conserved sites within the DENV genome of various subtypes, particularly in NS1, NS3, and NS5. Overexpression of miR-942-5p suppressed viral replication, establishing it as an antiviral miRNA. DENV infection suppressed endogenous miR-942-5p levels, and circEPSTI1 may further reduce its availability, thereby relieving repression on viral targets.

RNA-seq experiment identified SERPINE1 to be downstream of the circEPSTI1 and miR-942-5p, as knockdown of circEPSTI1 or overexpression of miR-942-5p suppressed the expression of SERPINE1. Moreover, reverse signature analysis revealed that the circEPSTI1 knockdown gene signature resembles that of the AKT1 knockout mouse. miR-942-5p target gene enrichment analysis also identified enrichment of the PI3K-AKT1 signaling pathway. SERPINE1 has been previously reported to activate the PI3K/AKT1 signaling pathway and influence cell death pathways in several cancers [15]. The AKT1 signaling activation is mediated through the phosphorylation of AKT1 at Serine 473 residue (p-AKT1). Hence, we speculated that the circEPSTI1-miR-942-5p-SERPINE1 axis regulates the AKT1 signaling pathway.

Multiple viruses utilize this pathway to promote their replication [16]. However, the interplay between AKT1 signaling and virus immunity is complex and has been reported to be both proviral and antiviral in different virus infections. Upon DENV infection, the level of p-AKT1 increased, indicating the activation of AKT1 signaling. SERPINE1 activates this pathway to enhance DENV survival. CircEPSTI1 knockdown or miR-942-5p overexpression reduced the p-AKT1 levels and suppressed the viral load. Hence, circEPSTI1-miR-942-5p-SERPINE1-AKT1 signaling axis regulates dengue infection and is a potential target for antiviral therapy. SERPINE1 inhibitor Tiplaxtinin was utilized to target this axis, and it not only decreased the viral load but also reduced the expression of circEPSTI1.

Overall, these findings reveal circEPSTI1 as a diagnostically and therapeutically relevant circular RNA in dengue infection. Through the identification of circEPSTI1-miR-942-5p-SERPINE1-AKT1 signaling axis, this study expands our understanding of circRNAs in the host-pathogen interactions and positions them as a crucial regulator in viral innate immune evasion. The findings also suggest the possibility that many other RNA viruses exploit similar circRNA-miRNA signaling pathway axes for their survival and could be new therapeutic targets for antiviral treatment. Additionally, the circEPSTI1 could be a significant diagnostic and prognostic marker, as its expression levels change with disease stages **(Figure 7)**.

This study also has limitations. Although circEPSTI1 induction upon DENV infection is conserved in mouse PBMCs ex vivo, establishing an in vivo model remains challenging since mice are not the natural hosts for the dengue virus. Moreover, the use of interferon receptor knockout mice complicates the system, as circEPSTI1 induction is interferon-dependent and therefore cannot be properly assessed under such conditions. Nevertheless, the conserved induction of circEPSTI1 across species highlights its potential role as a fundamental host response factor to viral infection. Future studies employing humanized mouse models or alternative primate systems may provide deeper insights into the circEPSTI1–SERPINE1–AKT1 regulatory axis during DENV infection. Importantly, the ability of Tiplaxtinin to suppress circEPSTI1 expression and reduce viral burden underscores the therapeutic potential of targeting this pathway, not only in dengue but possibly in other viral infections that exploit AKT1 signaling for replication. Future studies exploring the functional mechanism by which TPX regulate circEPSTI1 upon different pathological conditions may give insights to the therapeutic potential.

## Materials and Methods

### Patient samples

The Plasma samples utilized in the study were collected from the College of Medicine and J.N.M. Hospital, Kalyani, West Bengal, and Sanjay Gandhi Post-graduate Institute of Medical Science (SGPI), Lucknow, between January 2023 and April 2025.

Patients presenting to the hospitals with acute febrile illnesses were screened for dengue. The dengue diagnosis was performed either by the NS1-antigen test, IgM antibody test, or RT-PCR. The inclusion criteria include patients with clinically-confirmed dengue with either of the above tests, symptoms onset within 5 days before the sample collection, willingness to participate in this study through informed consent, and, in case of healthy volunteers, subjects with similar geographical origin, no recent history of acute febrile illnesses, and age and sex matched individuals compared to the dengue subjects. The exclusion criteria include pregnant women, chronic immunosuppression, confirmed cases of malaria or other febrile illnesses, low quality of clinical samples (haemolysis), and low yield of RNA. The age, sex, blood platelet counts, and symptoms were recorded for the study **(Supplementary Table S1)**.

### Human PBMC collection

The whole blood of volumes 3-5ml was collected from the clinical subjects in EDTA-coated blood collection tubes. Plasma was separated within an hour by centrifuging at 1500g for 10 minutes at room temperature and was stored at −80°C till RNA extraction. Human PBMCs were collected from healthy donors using EDTA blood tubes and isolated using Histopaque-1077 (Sigma) as per the manufacturer’s guidelines.

### Mice Experiments and Ethics Statement

The design and protocols for animal (mice) experiments were reviewed and approved by the Institute Animal Ethics Committee and the Institute Biosafety Committee (IBSC/IISERB/2025/Meeting-I/05). Three to five-week-old C57BL/6 mice were housed and bred under specific pathogen-free conditions in the Animal facility, IISER Bhopal. Blood (600 μL) was drawn from the mice using cardiac puncture. PBMC was isolated using Histopaque-1077 (Sigma) as per the manufacturer’s guidelines.

### Reanalysis of publicly available data from the NCBI-GEO database

Publicly available RNA-sequencing datasets related to Dengue virus infection clinical samples in which Total RNA-sequencing was performed were retrieved from the NCBI-GEO database. Only one dataset, GSE94892 was found eligible for the study [17, 18]. The raw sequencing reads were downloaded and quality-checked. Adapter trimming and low-quality read filtering were performed using fastp [19]. The clean reads were aligned to the human reference genome (GRCh38) using the BWA aligner [20]. The unmapped reads were further processed for circRNA detection using the CIRI2 algorithm [21]. The quantification of circRNAs was performed through CIRIquant [22]. The circRNA expression counts were pre-filtered with rowsums more than 5 in at least 10 samples to avoid the least-expressing circRNAs with zero counts. Differential expression analysis between DENV-infected and control samples was performed using DESeq2 [23]. The top differently expressed circRNAs (DEcircRNAs) were identified based on the adjusted p-value and the logFC. The top DECircRNAs were visualized through a heatmap, a violin plot, and a dot plot using the ggplot2 R package.

### Cells and transfection

A549 human alveolar basal epithelial cells (ATCC CCL-185), HEK293T human embryonic kidney cells (ATCC CRL-3216), HepG2 human hepatoblastoma cell line (HB-8065), and MRC5 Medical Research Council cell strain 5 (ATCC CCL-171) were grown in Dulbecco’s modified Eagle’s medium (DMEM) supplemented with 10% fetal bovine serum (FBS) and 1% penicillin-streptomycin. DNA, siRNA transfection was carried out using Lipofectamine 3000 (Invitrogen). For electroporation of human PBMCs, 1 × 10^6 cells were resuspended in Opti-MEM (Invitrogen) containing 50 nM miRNA mimics (ThermoFisher Scientific). The cells were subjected to two pulses of 1000 V for 0.5 ms with a 5-second interval between pulses using the Gene Pulser Xcell electroporation system. Following electroporation, the cells were transferred to RPMI supplemented with 10% FBS.

### Generation of the DENV2 virus

Dengue virus (strain DENV2, New Guinea C) was used in this study. Dengue virus was rescued from the reverse genetic system gifted by Prof. Andrew D Davidson, University of Bristol [24]. Briefly, the plasmid was cloned into a low-copy-number plasmid, pWSK29. Transformation of the vector pWSK29 was done into E. coli (DH5α) followed by overnight incubation and plasmid DNA isolation. Xba I linearized pDVWS601 plasmid was transcribed in vitro using a T7 transcription kit, followed by determination of the concentration of the RNA. For virus recovery, BHK-21 cells were transfected using DENV2 transcripts, and the virus was harvested 48 hours after transfection. The cells were fixed, over-dried, and then assayed for evidence of viral replication by immunofluorescence assay using an antibody recognizing the DENV NS5 protein. The virus was passaged further in C6/36 cells.

### Viruses and infection

Cells were washed with PBS and infected with 0.5 MOI DENV2 virus dissolved in SF-DMEM. After 1.5 hrs, cells were washed with PBS again and incubated with SF-DMEM supplemented with 1% FBS.

### Quantitative real-time reverse transcription-PCR

Intracellular RNA was extracted using TRIzol reagent (Invitrogen) while RNA from supernatant and patient samples were isolated by TRIzol LS reagent (Invitrogen) and then converted into cDNA with the High-Capacity cDNA synthesis kit (ThermoFisher Scientific) as per the manufacturer’s guidelines. Gene expression was evaluated through quantitative real-time PCR using gene-specific primers and PowerUp SYBR Green Master Mix (ThermoFisher Scientific) **(Supplementary Table S2-S3)**.

### RNA Fluorescence in situ hybridization

A549 cells were seeded in a 24-well plate and infected with DENV virus at an MOI of 0.5 for 36h. The cells were fixed with 4% formaldehyde for 20min and permeabilized with 0.1% Triton X-100 for 5 min. The cells were treated with prehybridization buffer (2X saline sodium citrate buffer, 10% formamide) for 20min at room temperature. The cells were then hybridized in hybridization buffer (10% Dextran sulfate, 4X SSC, 40% formamide, 10 × 10^−3^ M DDT, 100 µg/ml yeast tRNA, 100 µg/ml sheared salmon sperm DNA, 1X Denhardt’s solution) with 2 µl of Cy3-labeled circEPSTI1 probe at 55°C overnight. The cells were washed with PBS, and the nuclei were stained with DAPI. The images were taken with an LSM 780 confocal laser microscope.

### RNA-seq library preparation and high-throughput sequencing

The RNA Samples that passed the Quality control (QC) for concentration and integrity were subjected to ribosomal RNA depletion using the KAPA RiboErase HMR kit (Roche), followed by cDNA synthesis and amplification with the KAPA HyperPrep Kit (KK8544, Roche) and purification with KAPA pure beads (KK8545, Roche) to prepare the sequencing libraries. The library size distribution and assessment were performed using Agilent Bioanalyzer. The QC-passed paired-end libraries were then sequenced using the Illumina Novaseq 6000 instrument by Nucleome Informatics Pvt. Ltd, Hyderabad, India.

### RNA-seq data analysis to identify the significant genes

The raw reads were subjected to quality assessment and adapter trimming using the fastp tool [19]. The cleaned reads were then aligned to the human genome (hg19) using HISAT2 [25]. The mapped binary alignment files were quantified using Stringtie to obtain the raw counts. Normalization and differential expression analysis were performed using the R package DESeq2 [23]. The significant genes were identified using the threshold |logFC| >1 and adj.P.Val < 0.05. Enrichment analysis and the reverse gene signature analysis were performed using the EnrichR tool [26].

### Digital PCR

RNA was extracted with Trizol and cDNA synthesis was done using High-Capacity cDNA synthesis kit (ThermoFisher Scientific) as per the manufacturer’s guidelines. dPCR reactions were performed using the A549 cell line cDNA as template and circEPSTI1 primer using the kit QIAcuity EG PCR Kit (250111, QIAGEN).

### Luciferase reporter assay

HEK 293T cells were seeded in 24-well plates and transfected at 60–70% confluency with 30 ng of the control plasmid pRL-TK, 100 ng of the luciferase reporter plasmid, and 25 nM of the control or miR-942 mimic. Cells were collected and lysed 36 hours post-transfection, and luciferase activity was quantified using the Dual-Luciferase® Reporter Assay System (Promega, Cat# E1960) according to the manufacturer’s instructions, and the readings were recorded in the GloMax® Multi+ system (Promega).

### RNA immunoprecipitation

HEK293T cells were lysed using an ice-cold lysis buffer supplemented with 1X protease inhibitor cocktail (11836145001, Roche). The lysate was incubated with FLAG M2 affinity beads (Sigma) overnight at 4°C. The lysate was washed with washing buffer (300 mM NaCl, 50 mM tris-glycine (pH 7.5), 5 mM MgCl2, and 0.05% NP-40). The RNA was extracted using TRIZOL reagent.

### Biotin Pull-Down Assay

A549 cells were transfected with the empty vector and circEPSTI1 plasmid. The cells were lysed 24 hours after transfection and were incubated with control and circEPSTI1 biotin probe overnight at 4°C. The lysate was incubated with Pierce™ Streptavidin Magnetic Beads (88817, ThermoFisher Scientific) at 4°C for 2 hours. The beads were then washed thrice with lysis buffer, and the RNA was extracted using TRIZOL.

### Enzyme-linked immunosorbent assay

Culture supernatants from A549 cell lines and PBMC were collected and analyzed by Enzyme-linked immunosorbent assay following the manufacturer’s protocols to determine the levels of IP-10 (550926, BD OptEIA™ Human IP-10 ELISA Set) and IL-6 (555220, BD OptEIA™ Human IL-6 ELISA Set).

### Immunoblotting analysis

Cells were harvested 36 hours post-infection, washed with PBS, and lysed with ice-cold lysis buffer containing 1x protease inhibitor cocktail (11836145001, Roche) and 1mM of sodium orthovanadate (S6508, Sigma-Aldrich). The lysate was centrifuged at 15000g for 10 minutes at 4°C, and the supernatant was collected. Protein was quantified by Bradford assay, and around 10 µg of total protein was loaded. Immunoblotting was done using anti-AKT1 (9Q7, Invitrogen), anti-phospho-Akt (9271, Ser473, Cell Signaling Technology), anti-NS5 (MA5-17295, Invitrogen), anti-NS1 (SAB2700022, Sigma-Aldrich), anti-β actin (A1978, Sigma-Aldrich) antibody, anti-SERPINE1 (MA5-17171), anti-FLAG (F1804, Sigma-Aldrich). Secondary anti-mouse IgG and anti-rabbit IgG antibodies were purchased from Invitrogen. The immunoblotted PVDF membrane was visualized with the LI-COR system.

### Confocal microscopy

A549 cells were grown under cover slips in 12 well plate. The cells were fixed with acetone: methanol in 1:1 ratio for 10 mins at −20°C. The fixed cells were blocked with BSA supplemented with 0.1% Triton X-100 for permeabilization for 2 hours at room temperature and incubated with the primary antibody for 1 hour. The cells were washed three times with PBST and incubated with the secondary antibody for 30 minutes at room temperature. The cells were washed three times with PBST, the nucleus was stained with DAPI, and the coverslip was mounted on a glass slide. The cells were then analyzed with the LSM 780 confocal laser microscope. ImageJ software was used to analyse the images.

### Flow cytometry

Cells were harvested post-36 hours of infection, trypsinized, and fixed with 4% formaldehyde for 5 minutes. The cells were permeabilized with 0.1% Triton X-100 for 10 minutes, blocked with 5% FBS for 40 minutes, incubated with primary antibodies for 40 minutes, and secondary antibody for 30 minutes. The cells were analyzed using a flow cytometer, FACSAria III (Becton Dickinson), and the data were analyzed using FlowJo software.

### Statistical analysis

All experiments were conducted with appropriate control or mock-transfected samples. Each experiment was repeated independently two or three times. Data were analyzed for statistical significance using GraphPad Prism software. Differences between two groups were assessed using an unpaired, two-tailed Student’s t-test, while differences between three or more groups were evaluated using ANOVA followed by the Newman-Keuls test. Differences were regarded as statistically significant when P < 0.05. Statistical significance in the figures is represented as follows: ***P < 0.001, **P < 0.01, *P < 0.05; ns, not significant.

## Data availability

The RNA-seq data performed in the study have been submitted to the Gene Expression Omnibus under the accession number GSE306475. The reviewers can access the raw data using the secure token ylulcucipbuzxkx. The dataset will be made publicly available after the publication. All the data reported in this article are available in the main figures and the supplementary materials. Any further information can be requested from the corresponding author through.

## Acknowledgements

This work was supported by the intramural research grant from the Indian Institute of Science Education and Research, Bhopal (IISER Bhopal). The funders had no role in study design, data collection, interpretation, or the decision to submit the work for publication. The pIRESneo-Flag/HA Ago2 plasmid was a gift from T. Tuschl (Addgene; plasmid 10822). mc2 mNeon plasmid for circRNA overexpression was a gift from Simon Conn & Brett Stringer (Addgene plasmid # 206218). The DENV2 reverse genetics system was a kind gift from Prof. Davidson. We thank IISER Bhopal for providing access to the Central Instrumentation Facility.

## Contributions

**Nilanjana Das** – Conceptualization, Methodology, Validation, Formal Analysis, Investigation, Data curation, Writing – Original Draft, Writing-Review & Editing, Visualization.

**Pandikannan Krishnamoorthy** - Conceptualization, Methodology, Software, Formal Analysis, Investigation, Data curation, Writing – Original Draft, Writing- Review & Editing, Visualization.

**Kushal Ramrao Junghare-** Investigation, Validation, Visualization, Writing- Review & Editing.

**Athira S Raj-** Investigation, Visualization, Data curation, Writing- Review & Editing.

**Priyanka De**- Resources, Writing- Review & Editing

**Shouvanik Adhya -** Resources, Writing- Review & Editing

**Atul Garg –** Resources, Writing- Review & Editing

**Dheeraj Kheten –** Resources, Writing- Review & Editing

**Ashok Kumar-** Resources, Writing- Review & Editing

**Himanshu Kumar-** Conceptualization, Methodology, Formal Analysis, Data curation, Writing – Original Draft, Writing- Review & Editing, Project administration, Funding Acquisition Visualization, Supervision

## Ethics declarations

All the participants provided informed consent, and the study was approved by the Institutional Ethics Committee of COMJNMH, Kalyani (F-24/PR/COMJNMH/IEC/24/2/42), the Institutional Ethics Committee of SGPI, Lucknow (PGI/BE/338/2023), and the Institutional Ethical Committee of IISER Bhopal (IISERB/IEC/Certificate/20l7-II/01). The IBSC approval number for mice experiment is IBSC/IISERB/2025/Meeting-I/05.

## Supplementary Figure Legends

**Supplementary Figure 1:** (A) Principal Component Analysis (PCA) plot of circRNA expression profiles from PBMC samples. Control (blue) and DENV-infected (red) samples form distinct clusters based on variance along PC1 (26%) and РC2 (17%). Ellipses denote the 95% confidence intervals for each group. (B) Violin plot showing the distribution of log2-transformed expression values for the top five differentially expressed circRNAs between control and DENV-infected samples. Each point represents an individual sample. (C) Heatmap representing z-score normalized expression of the top 10 differentially expressed circRNAs across all samples. Red color indicates high expression, and blue color indicates low expression. Sample groupings (Control vs DENV) are indicated at the top of the heatmap.

**Supplementary Figure 2:** (A) Schematic illustration depicting the genomic locus and biogenesis of circEPSTI1. The circEPSTI1 is generated from the pre-mRNA of EPSTI1 by back-splicing between Exon 6(splice donor site) and Exon 2(splice acceptor site) leading to an exonic circular RNA of 375 bp. (B) circEPSTI1 and linear EPSTII1 levels were measured by qPCR in A549 and PBMC cells using cDNA synthesized from oligo-dT primers or random hexamer method and genomic DNA or cDNA (random hexamer method) as emplates. GAPDH was used as a positive control. (C) A549 and PBMC cells were pre-treated with Actinomycin D (10ug/mL) for the indicated times, further, the expression of linear EPSTI1 and circEPSTI1 was tested by qPCR. (D) circEPSTI1 was quantified in RNA isolated from the cytoplasmic and nuclear cell fractionation of DENV-infected (48hr) A549 cell line. GAPDH and U6 were used as the positive controls for the cytoplasmic and nuclear fractions respectively. (E) DENV infected (48hr) A549 cellular RNA was treated with and without RNase R (2U/uL), and the linearEPSTI1 and circularEPSTI1 were quantified by qPCR.

**Supplementary Figure 3:** (A) Human PBMCs (n=11) were isolated and infected with DENV2 at MOI=2 for 36hrs. The viral load was quantified by qPCR. (B) Conservation of human and mouse circEPSTI1. (C) Mouse PBMCs (n=3) were isolated and infected with DENV2 at MOI=4 for 36hrs. The viral load was quantified by qPCR. (D) HepG2 and MRC5 cell lines were infected with DENV at an MOI=1 respectively for the indicated time points and circEPSTI1 was quantified by qPCR. (E) PBMCs, A549, HepG2 and MRC5 cell lines were infected with DENV at an MOI=0.5 for the indicated time points and the viral load was quantified by qPCR. (F) A549, MRC5 and PBMC cells were transfected with 25nM of circEPSTI1 siRNA and infected with DENV for 36h. The levels of linear EPSTI1 were quantified by qPCR.

**Supplementary Figure 4:** (A) RNA immunoprecipitation of AGO2 was done and the levels of miR-942-5p was quantified by qPCR in HEK293T cells. (B) HEK293T cells were transfected with 2µg of plasmid encoding FLAG-AGO2 and 25nM mir-942 and were infected with DENV virus (MOI=2) for 36h. AGO2 pulldown was checked by western blot using FLAG antibody. (C) EK293T cells were infected with DENV for 36 hrs. A biotin-labeled circEPSTI1 probe was used to pull down the RNA fractions binding to the circEPSTI1, and the level of circEPSTI1 was quantified by qPCR.

**Supplementary Figure 5:** (A) HEK293T cells were transfected with 2µg of plasmid encoding FLAG-AGO2 and 25nM mir-942 and were infected with DENV virus (MOI=2) for 36h. The levels of DENV were quantified by qPCR. (B) A549 cells were transfected with 25nM of miR-942-5p mimic for 24h. The levels of miR-942-5p was quantified by qPCR. (C) A549 cells were transfected with 25nM of miR-942 inhibitor for 24h. The levels of miR-942-5p was quantified by qPCR. (D) A549 cells were transfected with 25nM of miR-942-5p mimic for 24hr and infected with DENV for 36h. The viral load was quantified by qPCR in the supernatant. (E) A549 cells were transfected with 25nM of sicircEPSTI1 and 25nM of miR-942 inhibitor simultaneously for 24hr and infected with DENV for 36h. The levels of miR-942-5p and DENV were quantified by qPCR.

**Supplementary Figure 6:** (A) Expression plots showing the expression of the common genes between circEPSTI1 knockdown and miR-942-5p targets. (B) A549 cells were infected with DENV, and the level of SERPINE1 was quantified by qPCR at different time points by qPCR. (C) A549 cells were treated with (0.5ug/ml) recombinant IFN-β, and the level of circEPSTI1 was quantified at different time points by qPCR.

**Supplementary Figure 7:** (A) 549 cells were transfected with 25nM of siAKT1 and infected with DENV2 for 36h. The levels of IFN-β and OAS-2 were quantified by qPCR. (B) The A549 cells were pre-treated with AKT1 inhibitor Wortmannin and infected with DENV for 36h. The levels of DENV NS5, p-AKT1 and β-actin proteins were quantified by western blot. (C) The levels of circEPSTI1 were quantified by qPCR in Wortmannin-treated A549 DENV2 infected cells. PBMC cells were pre-treated with 5,10 and 20μM of Tiplaxtinin, a SERPINE1 inhibitor, for 2 hours, followed by infection with DENV (MOI=0.5) for 36h. The levels of DENV2 (D), (E) circEPSTI1 and (F) linear EPSTI1 were quantified by qPCR.

## Supplementary Table Legends

**Supplementary Table S1:**

Clinical and metadata information of the Indian dengue cohorts.

**Supplementary Table S2:**

List of Primers for Cloning and SDM.

**Supplementary Table S3**:

List of Primers for RT-PCR.

