## Supplementary figures and images for "CircEPSTI1 regulates miR-942-5p-SERPINE1-AKT1 signaling axis to enhance dengue infection and is suppressed by Tiplaxtinin"

### Supplementary Figures

**Supplementary Figure 1**

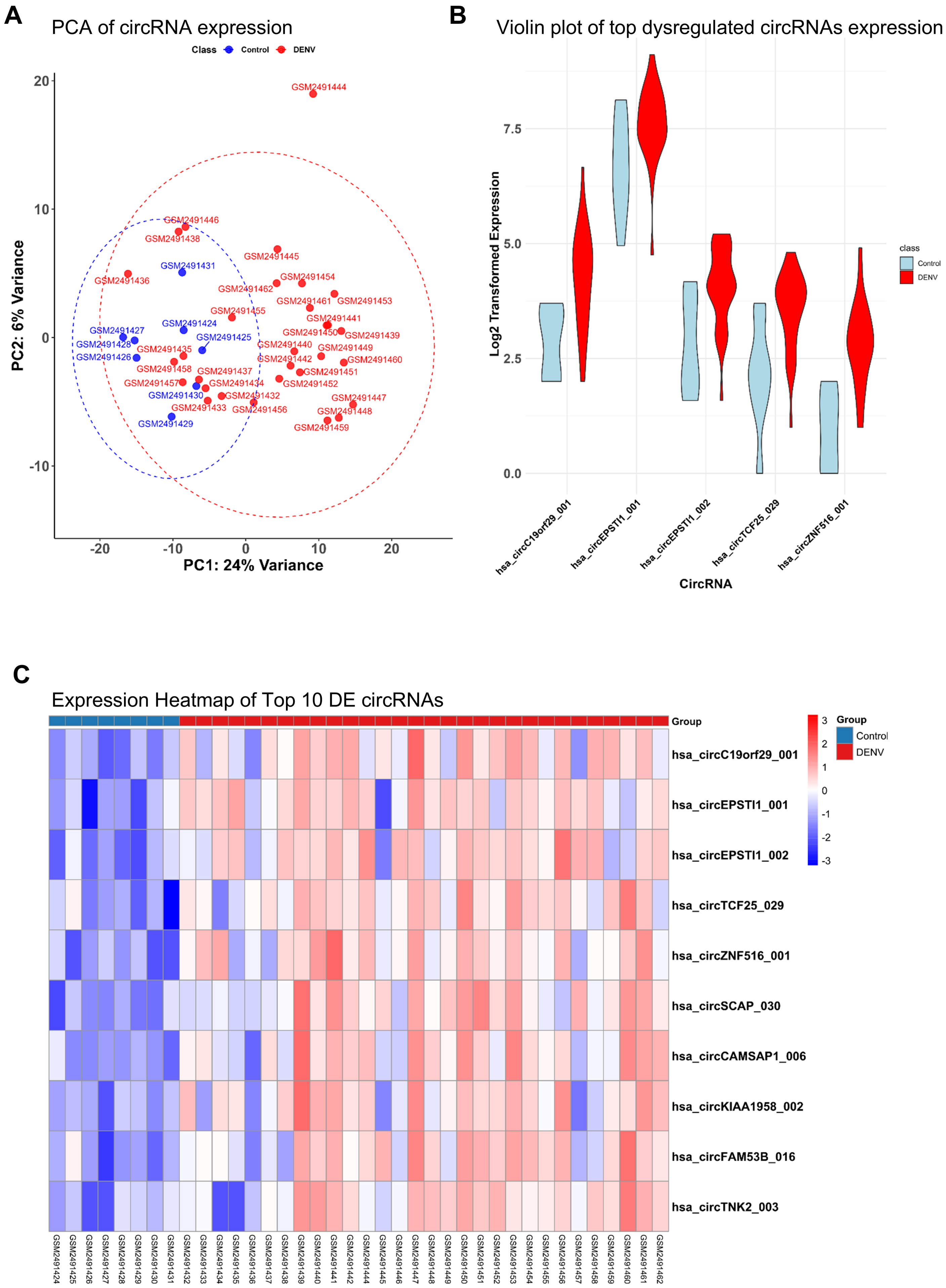

##### Supplementary Figure 2

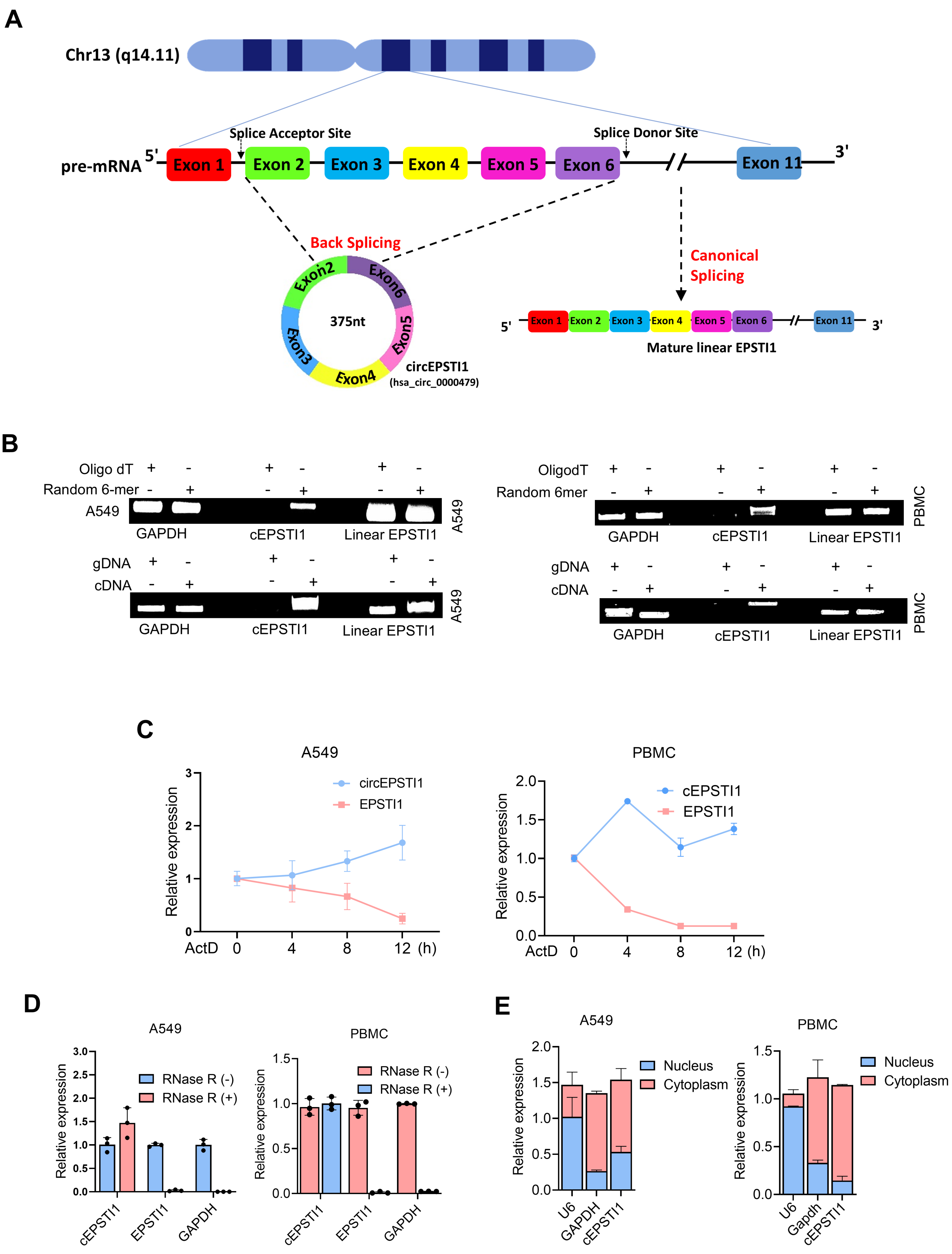

##### Supplementary Figure 3

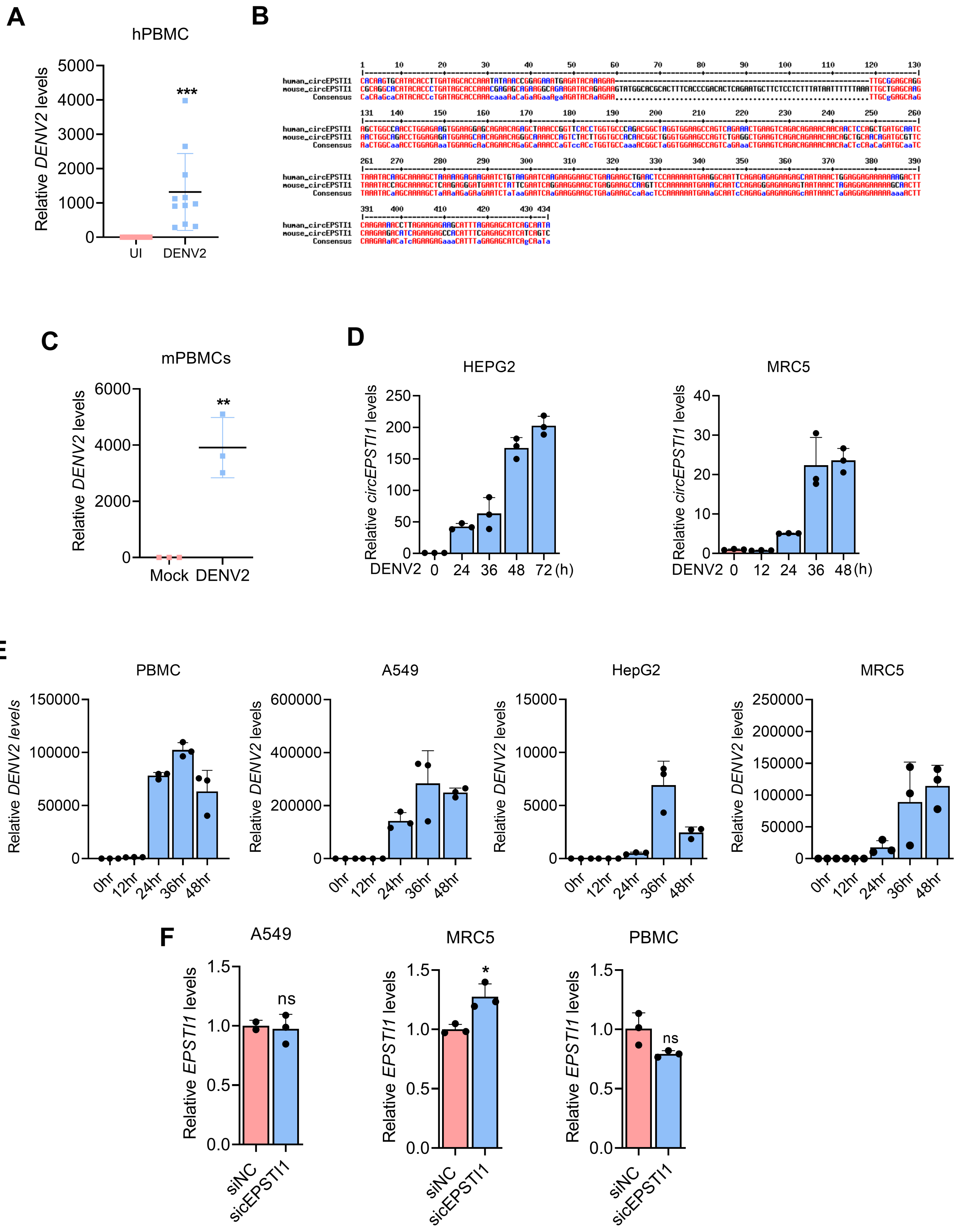

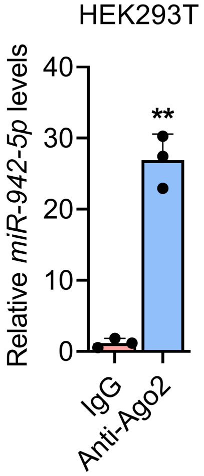

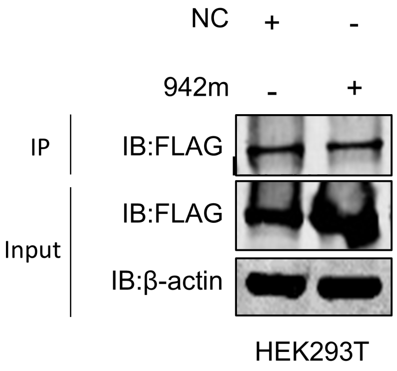

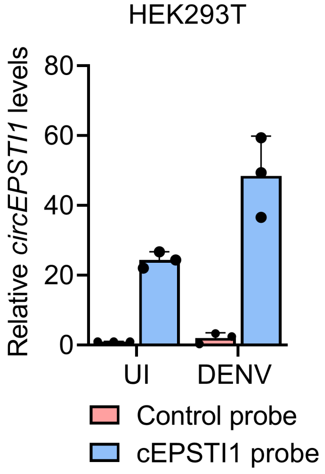

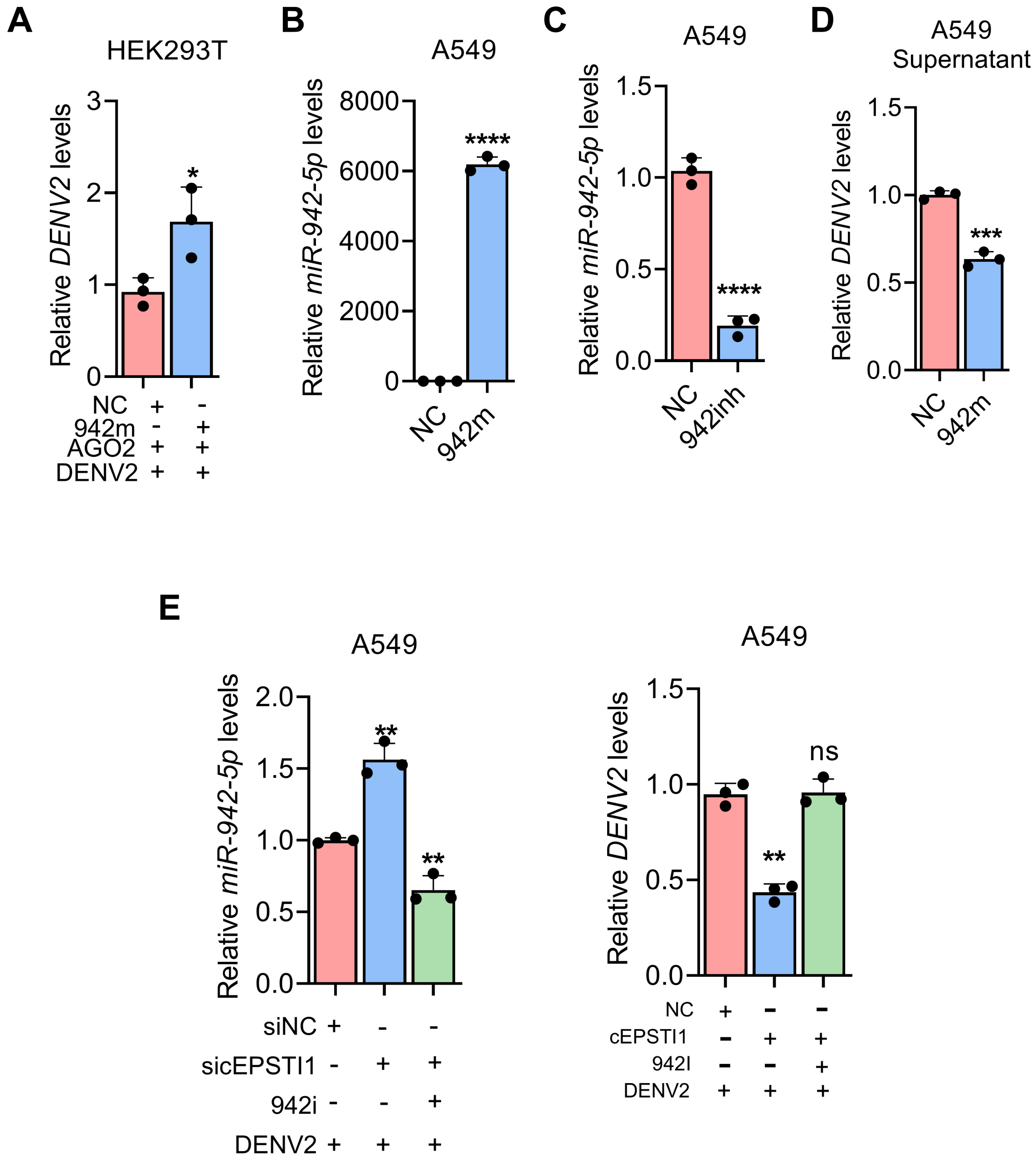

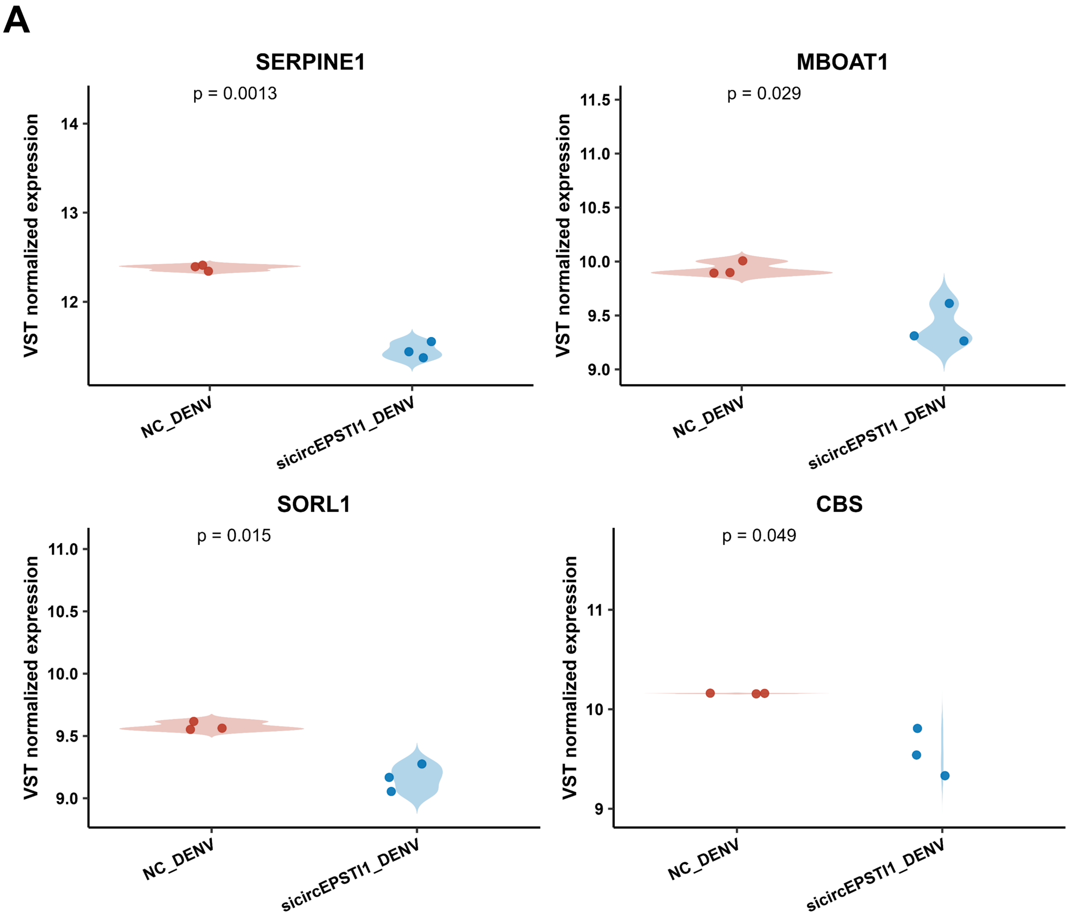

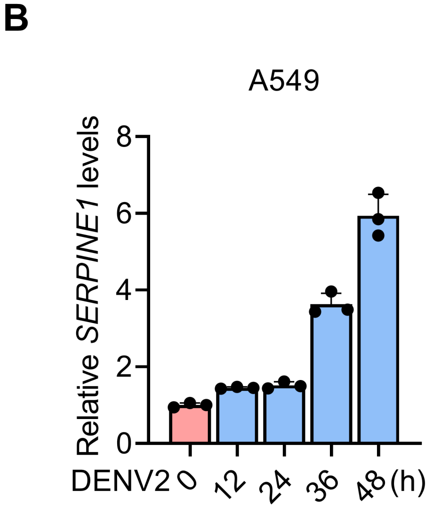

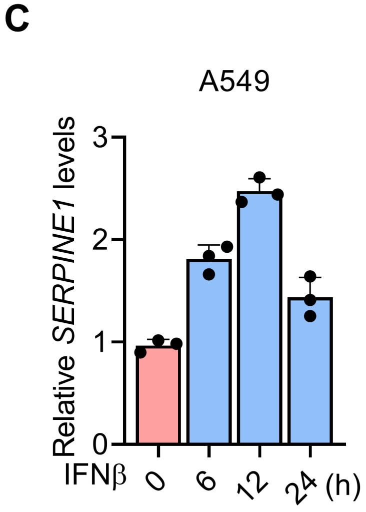

#### Supplementary Figure 7

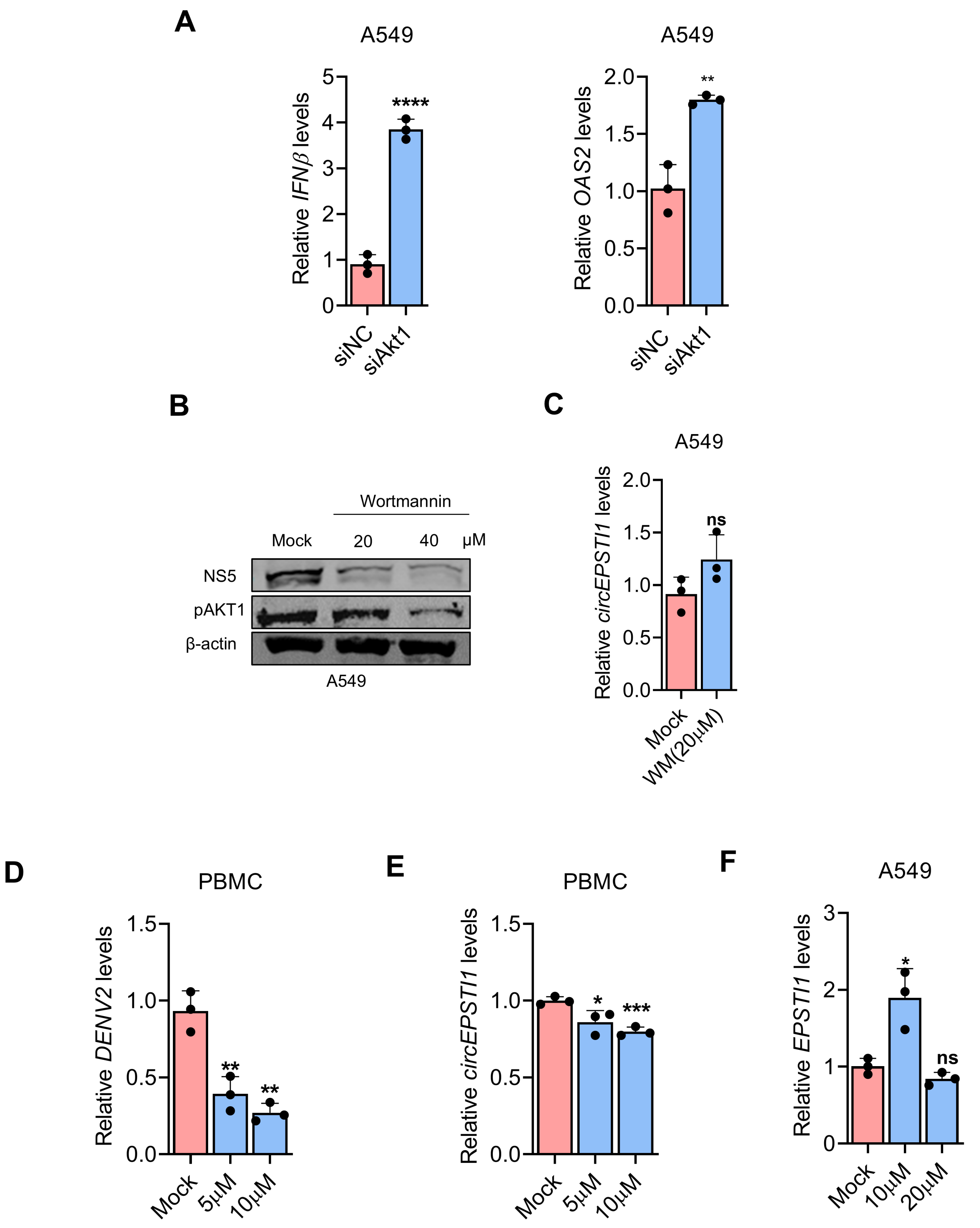
