## Supplementary Tables for "CircEPSTI1 regulates miR-942-5p-SERPINE1-AKT1 signaling axis to enhance dengue infection and is suppressed by Tiplaxtinin"

**Supplementary Table S1**:

### Clinical and metadata information of the Indian dengue cohorts.

| Group | N | Sex (M/F) | Age (mean ± SD) | Age range | Platelet Count (mean ± SD) | Platelet range | Diagnosis Tests distribution |
|---|---|---|---|---|---|---|---|
| **Healthy** | 56 | 37 / 19 | 29.6 ± 7.8 | 21–57 | NA | NA | NA (all healthy controls) |
| **Dengue** | 51 | 35 / 16 | 27.5 ± 20.4 | 1–70 | 108,639 ± 64,865 /µL | 15,000–292,000 | IgM: 22; NS1: 11; RT-PCR: 18 |

**Supplementary Table S2**:

**List of Primers for Cloning and SDM***

| Gene | Primer Sequences |
|---|---|
| PAI-3'UTR_SpeI_Fw | TTAC ACTAGT CCTGGGGAAAGACGCCTT |
| PAI-3'UTR_HindIII_Rv | TTAC AAGCTT AGTGCCACAGTGGACTCT |
| PAI_BamHI_ Fw | TTAC GGATCC ATGCAGATGTCTCCAGCC |
| PAI_XhoI_Rv | TTAC CTCGAG TCAGGGTTCCATCACTTGG |
| AKT1-3'_UTR_SpeI_Fw | TTAC ACTAGT CGTGCCATGATCTGTATTTAATGG |
| AKT1-3'_UTR_HindIII_Rv | TTAC AAGCTT AGATGACAGATAGCTGGTGA |
| AKT1_BamHI_Fw | TTAC GGATCC ATGAGCGACGTGGCTATTGTG |
| AKT1_XhoI_Rv | TTAC CTCGAG TCAGGCCGTGCCGCTGGCCGAGTA |
| circEPSTI1_BsmBI_Fw | GCGTCTCA TCAG CACAAGTGCATACACCT |
| circEPSTI1_BsmBI_Rv | ACGTCTCG TTAC TATTGCTGATGCTCTCT |
| circEPSTI1_SpeI_Fw | TTAC ACTAGT CACAAGTGCATACACCTTGA |
| circEPSTI1_HindIII_Rv | TTAC AAGCTT TATTGCTGATGCTCTCTAAATG |
| DENV2_NS1_Spe1_Fw | TTAC ACTAGT GATAGTGGTTGCGTTGTGAGC |
| DENV2_NS1_Mlu1_Rv | TTAC ACGCGT GGCTGTGACCAAGGAGTTGAC |
| DENV2_NS3_Spe1_Fw | TTAC ACTAGT GCTGGAGTATTGTGGGATG |
| DENV2_NS3_HindIII_Rv | TTAC AAGCTT CTTTCTTCCAGCTGCAAACTC |
| DENV2_NS5_Spe1_Fw | TTAC ACTAGT GGAACTGGCAACATAGGAGAGAC |
| DENV2_NS5_HindIII_Rv | TTAC AAGCTT CCACAGGACTCCTGCCTCTTC |
| PAI-3'UTR_Mutant1_Fw | CTTTCTGAAGGAAGATGGGACATTTG |
| PAI-3'UTR_ Mutant1_Rv | CAAAAGGCAAATGTCCCATCTTCCTT |
| PAI-3'UTR_ Mutant2_Fw | TGAGACCCTGGGAGATGGGTTTGAAG |
| PAI-3'UTR_ Mutant2_Rv | AGTTGTGCTTCAAACCCATCTCCCAG |
| Akt1-3'UTR_Mutant_Fw | CGGGTGCATTTGAGATGGGCCACGCTGTCCTCT |
| Akt1-3'UTR_Mutant_Rv | AGAGGACAGCGTGGCCCATCTCAAATGCACCCG |
| circEPSTI1_Mutant_Fw (224-231) | CAAAAGCTAAAAAGATGGGAATCTG |
| circEPSTI1_Mutant_Rv(224-231) | TGATTCTTACAGATTCCCATCTTTTTA |
| circEPSTI1_Mutant_Fw (295-302) | GGCAATTCAGAGAGATGGGAGCAATA |
| circEPSTI1_Mutant_Rv(295-302) | TCCAGTTTATTGCTCCCATCTCTCTG |
| circEPSTI1_Mutant_Fw (347-353) | GAAAACCTTAGAAGATGGGCATTTAG |
| circEPSTI1_Mutant_Rv(347-353) | ATGCTCTCTAAATGCCCATCTTCTAA |

*SDM : Side-Directed Mutagenesis

**Supplementary Table S3**:

### List of Primers for RT-PCR

| Gene | Primer Sequences |
|---|---|
| SERPINE 1_Fw | TCATGCCCCACTTCTTCAGG |
| SERPINE1_Rv | CCACTGGCCGTTGAAGTAGA |
| Akt1_Fw | GCTCACCCAGTGACAACTCA |
| Akt1_Rv | CTTGGCCACGATGACTTCCT |
| circEPSTI1_Fw | GGCAATTCAGAGAGAGAAGAGC |
| circEPSTI1_Rv | CCTTCCACTTCTCCAGGTTG |
| Linear_EPSTI1_Fw | GACAGAAGTGCCTGTCAAAGTG |
| Linear_EPSTI1_Rv | GCCGTTTCAGTTCCAGTAATTC |
| DENV2_Fw | TTGAGTAAACTGTGCAGCCTGTAGCTC |
| DENV2_Rv | GGGTCTCCTCTAACCTCTAGTCCT |
| GAPDH_Fw | ATCATCCCTGCCTCTACTGG |
| GAPDH_Rv | GTCAGGTCCACCACTGACAC |
| U6_Fw | CTCGCTTCGGCAGCACAT |
| U6_Rv | TTTGCGTGTCATCCTTGCG |
| IFN-β_Fw | ACAGACTTACAGGTTACCTCCGAAC |
| IFN-β_Rv | CATCTGCTGGTTGAAGAATGCTT |
| OAS2_Fw | GCTCCTATGGACGGAAAACA |
| OAS2_Rv | CAAGGGACTTCTGGATCTCG |
| Mouse_circEPSTI1_Fw | TCGAGAGCATCATCAGTC |
| Mouse_circEPSTI1_Rv | TGCTCTCGTTTGGTGCTATC |
